# A *de novo* designed enzyme for photo-proximity labeling of E3 ligase neighborhoods in live cells

**DOI:** 10.64898/2026.07.22.739993

**Authors:** Nam Hyeong Kim, Zhi Lin, Sangwon Nam, Danielle L. Swaney, Nevan Krogan, Yong Ho Kim, Michael J. Therien, William F. DeGrado, James A. Wells

**Affiliations:** Department of Pharmaceutical Chemistry, University of California, San Francisco, San Francisco, California, USA; Cardiovascular Research Institute, University of California, San Francisco, San Francisco, California, USA; Department of Chemistry, Duke University, Durham, North Carolina, USA; J. David Gladstone Institutes, San Francisco, California, USA; Quantitative Biosciences Institute, University of California, San Francisco, San Francisco, California, USA; Department of Bioengineering and Therapeutic Sciences, University of California, San Francisco, San Francisco, California, USA; SKKU Advanced Institute of Nanotechnology (SAINT), Sungkyunkwan University (SKKU), Suwon, Gyeonggi-do, Republic of Korea; Department of Nano Science and Technology, Sungkyunkwan University (SKKU), Suwon, Gyeonggi-do, Republic of Korea; Institute of Multidisciplinary Research for Advanced Materials, Tohoku University, Sendai, Japan; Department of Cellular and Molecular Pharmacology, University of California, San Francisco, San Francisco, California, USA

**Author notes:** Corresponding authors: Zhi Lin, James A. Wells and William F. DeGrado. Designates co-equal first author (Nam Hyeong Kim, Zhi Lin). State Key Laboratory of Synergistic Chem-Bio Synthesis, School of Chemistry and Chemical Engineering, Shanghai Jiao Tong University; Shanghai Academy of Sciences (SANS), Shanghai, China.

**Keywords:** Photo-proximity labeling proteomics, *De novo* protein design, Domain-swapped dimer, Target identification, E3 ligase interactomes

## Abstract

Photocatalytic proximity labeling proteomics (photo-PLP) has emerged as a powerful technology for rapid capture of protein interactomes in situ. Typically, photo-PLP relies on chemical conjugation of the photocatalyst to the target of interest which creates practical challenges for derivatized photocatalyst synthesis and bioconjugation specificity. Integrating the precision of genetically encodable enzymes with the versatility of chemically defined photocatalysts provides a modular approach to further expand the scope of neighborhood mapping. Here, we present EYClamp, a *de novo* designed proximity labeling enzyme harnessing the off-the-shelf photocatalyst Eosin Y (EY) as a cofactor. Using a domain-swapped dimer architecture, we designed a scaffold that binds EY with high affinity (*K*_d_ = 10 nM) and lengthens its triplet excited-state lifetime by 29-fold. EYClamp enables efficient, multi-scale photocatalytic proximity labeling in live cells with aryl-diazirine-, aryl-azide- and phenol-biotin. We genetically fused EYClamp to a panel of six important E3 ligases. Using EYClamp, we identified over 1,500 candidate neighbors for KEAP1, MDM2, ASB7 and STUB1, providing a broad and unbiased view of these important neighborhoods. Critical functional networks were revealed including ASB7 engagement with HP1a/CUL5 complex for heterochromatin remodeling. Our EYClamp provides a genetically encodable “plug-and-play” solution for photo-PLP interactome discovery of the large family of E3 ligases and establishes domain-swapping as a promising strategy for *de novo* photoenzyme design.

## Introduction

Protein-protein interactions are critical regulators of cell biology by forming dynamic and transient interactome networks. Proximity labeling proteomics (PLP) has emerged as a powerful tool to study fragile protein neighborhoods in situ.^1–8^ Pioneering PLP systems such as BioID, TurboID, APEX2, NEDDylator and PUP-IT,^9–17^ were engineered from natural enzymes that are genetically fused to the protein of interest (POI), obviating the need for protein purification and conjugation. Proximity labeling is achieved through local chemical activation of biotinylated probes and shown to have broad utility in target identification campaigns amongst numerous biological scenarios.^18–24^

Recently, photocatalytic PLP (photo-PLP) platforms have been developed that covalently attach synthetic photocatalysts to the POI or POI-specific antibodies and ligands.^25–33^ When irradiated with visible light, the photocatalyst can trigger local activation of highly reactive biotinylated photo-probes and achieve interactome mapping with high temporal and spatial resolution.^34–36^ The recent MultiMap strategy used Eosin Y (EY) derivatives for multi-scale interactome profiling, where EY can activate different photo-probes with increasing labeling radii from aryl-diazirine-, aryl-azide-, to phenol-biotin, thus enabling neighborhood labeling at distinct length scales.^37–39^ In these applications, photo-PLP requires chemical modification of the photocatalyst and/or bioconjugation to the purified POI or antibody, which is more laborious than a genetically encoded system.^40–42^ Bridging genetically encoded protein scaffolds with precisely designed photocatalytic chemistry would simplify photo-PLP and broaden its utility.

To establish a more accessible and cost-effective platform for live-cell photo-PLP, we sought to develop a *de novo* protein^43–49^ capable of binding EY non-covalently. However, designing scaffolds for bulky, highly polar molecules like EY presents a formidable structural challenge. This difficulty arises from EY’s extended dimensions, the orthogonal arrangement of its xanthene and benzoic carboxylate rings, and its dianionic state at physiological pH resulting from its multiple polar groups. While previous *de novo* design efforts have generated small-molecule binders, these campaigns typically exhibit low success rates or are largely restricted to relatively apolar targets.^50–55^ Furthermore, although canonical four-helix bundles are the go-to solutions for many designed binders,^56–62^ the narrow, elongated cavities inherent to these folds cannot readily accommodate the bulky properties of EY. To overcome these limitations, we propose to engineer an inter-domain binding site by employing a domain-swapped dimer architecture. This strategy is widely utilized in nature and suggested to us that it might be an enabling method for *de novo* protein design.^63–65^

Here we present EYClamp, a genetically encodable photoenzyme for multi-scale proximity labeling in live cells. An important challenge was to tightly sequester the cofactor to enhance its photophysical properties, while providing sufficient accessibility for adequate reactivity with different photo-probes. Capitalizing on the structural versatility of domain-swapped dimers to accommodate complexes, asymmetric interfaces, we constructed a *de novo* protein domain that binds EY in a highly protected environment with high affinity (dissociation constant, *K*_d_ = 10 nM). Biophysical characterization confirmed that the precise structural design confined EY in a protected microenvironment, thereby extending its triplet excited-state lifetime by 29-fold. This triplet excited-state then transfers energy to activate probes such as aryl-diazirine-, aryl-azide- or phenol-biotin^37,38^ and form highly reactive carbenes, nitrenes or phenol radicals, respectively, for proximity labeling.

We demonstrate the cellular utility and remarkable specificity of EYClamp by fusing it to six important E3 ligases (KEAP1, MDM2, PARKIN, ASB7, RNF43, STUB1). E3 ligases are fate-determining regulators of cell biology through ubiquitination and protein turnover.^66^ However, the scope of E3 ligase substrates and their interactomes has been hampered by lack of high-resolution proximity labeling studies. Moreover, harnessing E3 ligases for intracellular targeted protein degradation has become a critical new modality in drug discovery,^67^ and expanding our understanding of their biology could help fuel that field. Using EYClamp for photo-PLP, we identified a total of 1633 neighbors for KEAP1, MDM2, STUB1 and ASB7, including neighbors such as HP1a (CBX5) and CUL5 that have recently been shown to engage with ASB7. Furthermore, by using the neighborhood identities, we successfully connected the associated functional networks of ASB7 with chromatin remodeling activity and STUB1 with cell-cell recognition. EYClamp can be expanded to other E3 ligases and more broadly as a simple, modular tool for photo-PLP and targeted interactome profiling. We also believe the domain-swapping design strategy can be applied to create other photocatalytic or co-factor dependent enzymes.

## Results

### Design principles for EYClamp

We sought several advantages over photo-PLP approaches that rely on covalent attachment of the photocatalyst to the POI or antibodies to it (**Figure 1A**): 1) a *de novo* EY-binder would use affordable, underivatized EY, enabling widespread adoption for labs without synthetic capabilities or the resources to purchase derivatives from specialty suppliers (**Supplementary Figure 1A**); 2) The lifetime and photophysical properties, which define reactivity and labeling efficiency, depend markedly on environment, which can be programed within a *de novo* binder. By contrast, solvent-exposed EY from our previously demonstrated antibody-based strategy^37^ and ecto-tag^38^ strategy limits the possibility for environmental fine tuning; 3) Tight, but reversible binding of EY can allow exchange of any photobleached EY protein complexes with unmodified EY in solution; 4) *De novo* proteins are extremely stable to both extreme pH and high temperature, making it possible for future applications with extremophiles. For example, a set of rhodamine binders were recently shown to enable imaging of a thermophile at extreme pH of 2 – 3 at a temperature reaching 75°C, which were not achievable with HaloTag and fluorescent proteins.^68^ Thus, we envision a *de novo* binder of EY could greatly extend the utility of photo-PLP for target identification.

**Figure 1:**
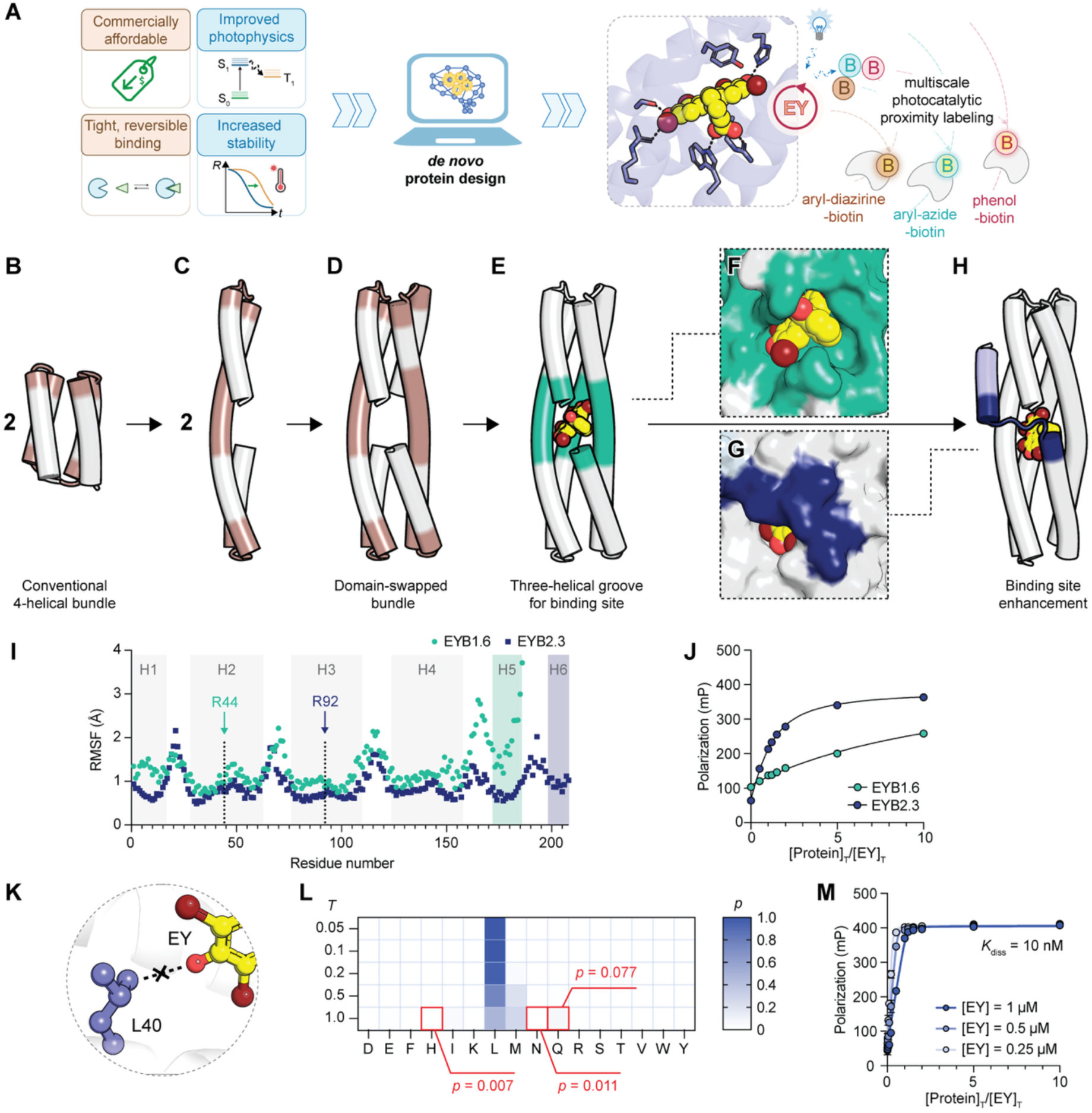
Design principles for an EY binder, EYClamp. (**A**) Rationale of designing an artificial proximity labeling enzyme using *de novo* protein design, which enables genetically encodable live cell multi-scale interactome profiling. (**B-H**) Schematic explaining the concept of domain-swapping (panels **B-D**), and the tailored design process (panels **E-H**). Starting from conventional 4-helical bundle (**B**), the loop interconnecting the second and third helices is converted into a monomer with a long extended helix (**C**), which can self-associate to form a domain-swapped dimer (**D**), as seen frequently in natural proteins. Our design began with this topology of a domain-swapped dimer and proceeded through designs that lifted the symmetry and enabled binding to EY (for detailed methods, see the ‘Overview of design process’ section in the **Supplementary Information**). The domain-swapped helices were interconnected by extending the helical strands to a length determined by the shape complementary to EY (**E**). Surface representations illustrate how the addition of a C-terminal helix extension (**H**) shields the solvent-exposed surface of bound EY (**F-G**). (**I**) Root-mean-square fluctuation (RMSF) analysis derived from unbiased all-atom molecular dynamics (MD) simulations of EYB1.6 and EYB2.3. H1–H6 denote the constituent helices of each candidate scaffold. (**J**) Binding affinity of EYB1.6 and EYB2.3 determined via a fluorescence polarization (FP) assay. The protein was titrated from 0 to 10 molar equivalents in the presence of 1 µM EY, with excitation at 460 nm (± 20 nm) and emission detected at 516 nm (± 10 nm). The fitting curves were calculated using the binding isotherm equation (for detailed methods, see the ‘Fluorescence emission and polarization measurement for EY binding assessment’ section in **Supplementary Information**). (**K**) Structural model illustrating the local microenvironment surrounding residue L40 and the 6-oxido atom of EY. (**L**) Temperature-dependent probability distribution for amino acid placement at the ketone-coordinating position, as predicted by LigandMPNN. **(M)** Binding affinity of EYClamp determined via FP assay. Polarization was recorded by titrating EYClamp (0 to 10 molar equivalents) against three initial concentrations of EY (0.25, 0.5, and 1 µM). The dissociation constant (*K*_d_) was derived from a global fit of the binding isotherms across all three EY concentrations (for detailed methods, see **Supplementary Information**).

To realize these advantages, we engineered an inter-domain binding site by employing a domain-swapped dimer architecture (**Supplementary Figure 1B**). Specifically, we deleted the loop between the second and third helices of a four-helix bundle (**Figure 1B-C**), converting previously intramolecular interactions into intermolecular contacts within the dimer (**Figure 1D**). This topological shift repurposes a region that formerly served only as a helical connector into an active structural interface for target molecule engagement. Augmented by the terminal ends of two adjacent helices, the groove formed by the three central helices provides a wider, more versatile binding pocket (**Figure 1E**). In this construct, the binding site is located precisely at the interface between the two swapped domains. Using a parametric model, we generated a family of these domain-swapped bundles to serve as binding scaffold (see the ‘Overview of design process’ section in the **Supplementary Methods**). We then utilized the COMBS^69^ method to determine the optimal orientation of EY within the scaffold and to identify amino acids capable of specifically coordinating the chemical moieties of the ligand. Specifically, we constrained an Arg residue to establish a strong bidentate electrostatic interaction between its guanidinium group and the EY carboxylate. The remaining binding residues were selected by evaluating candidate amino acids capable of acting as hydrogen bond donors to coordinate the ketone and hydroxyl groups of EY. Among the seven candidates experimentally tested (**Supplementary Data Table 1**), EYB1.6 exhibited the highest binding affinity for EY, with a *K*_d_ of approximately 10 μM (**Supplementary Figure 1C**).

All-atom MD simulations of EYB1.6 indicated that the C-terminal helix, which composes a portion of the binding site, was poorly organized with a backbone RMSF of 2 – 4 Å. Moreover, the solvent-exposed interaction between the EY carboxylate and R44 was not retained in an ideal geometry over the course of the simulation. Indeed, Ala mutations in the binding site were consistent with the MD simulations and showed that R44 did not significantly contribute to binding (**Supplementary Figure 1D**). To address this problem, we built a 20 amino-acid C-terminal extension to better enclose the ligand and to enhance the structural stability of the binding site (**Figure 1E-H, Supplementary Figure 1E**). This extension forms a short loop that drapes across the upper part of the binding site (as viewed in **Figure 1F and G**) followed by a 15-residue helix, which inserts into the N-terminal bundle, to create a 5-helix bundle. We also re-oriented the EY molecule to allow it to form a more well-shielded interaction with R92 deeper in the binding pocket. Unbiased all-atom MD simulations confirmed that this extension dampened backbone fluctuations, particularly within the H5 strand, therefore enabling stable EY binding (**Figure 1I, Supplementary Figure 1F**). The resulting design, EYB2.3, achieved a significantly improved *K*_d_ of 0.64 µM (**Figure 1J** and **Supplementary Figure 1G**).

Interestingly, LigandMPNN struggled to sample polar donors at position 40, assigning exceptionally low probabilities to His, Asn, and Gln even at elevated sampling temperatures (**Figure 1K-L** and **Supplementary Figure 1H**). This limitation highlights training distribution bias associated with the limited size of training data for protein complexes with synthetic small molecules, particularly with the infrequently encountered substitution of multiple Br atoms adjacent to an oxido group within the tricyclic ring. To bridge this gap, we employed structure-guided rational design. Targeted substitution of L40 with these polar residues dramatically improved the binding affinity, demonstrating that human reasoning and physics-based models can increase affinity when applied to microenvironments that are not frequently encountered in the artificial intelligence (AI) training sets (**Figure 1L**). This mutation resulted in a 10 to 100-fold improvement in *K*_d_; titrating protein into a fixed concentration of EY yielded a linear increase in fluorescence with an abrupt leveling at a stoichiometry of precisely 1, indicating a 1:1 ratio tight binding affinity. A sensitivity analysis showed *K*_d_ was < 30 nM, and *K*_d_ = 10 nM was obtained from non-linear least squares analysis (**Figure 1M**, see **Supplementary method**).

### Characterization of the binding and biophysical properties of EYClamp

The UV-Vis absorption and fluorescence emission spectra of EY exhibited pronounced shifts upon binding to EY, reflecting its incorporation into the structurally confined protein microenvironment. During protein titration, the primary absorbance peaked at 518 nm decreased, accompanied by the emergence of a new bathochromic band at 525 nm (**Figure 2A**). Additionally, while the fluorescence intensity of EY initially decreased with a slight red shift, increasing EYClamp concentration led to the fluorescence recovery with a new emission band emerged at 539 nm (**Figure 2B**). Circular dichroism (CD) spectroscopy confirmed that EYClamp maintains a well-folded α-helical structure and exhibits high thermal stability in melting experiments (**Figure 2C** and **Supplementary Figure 2A-B**). Furthermore, although EY is achiral and hence does not show optical activity, it shows a strong circular dichroic spectrum when it is rigidly bound in the chiral EY binding site (**Figure 2D**).

**Figure 2:**
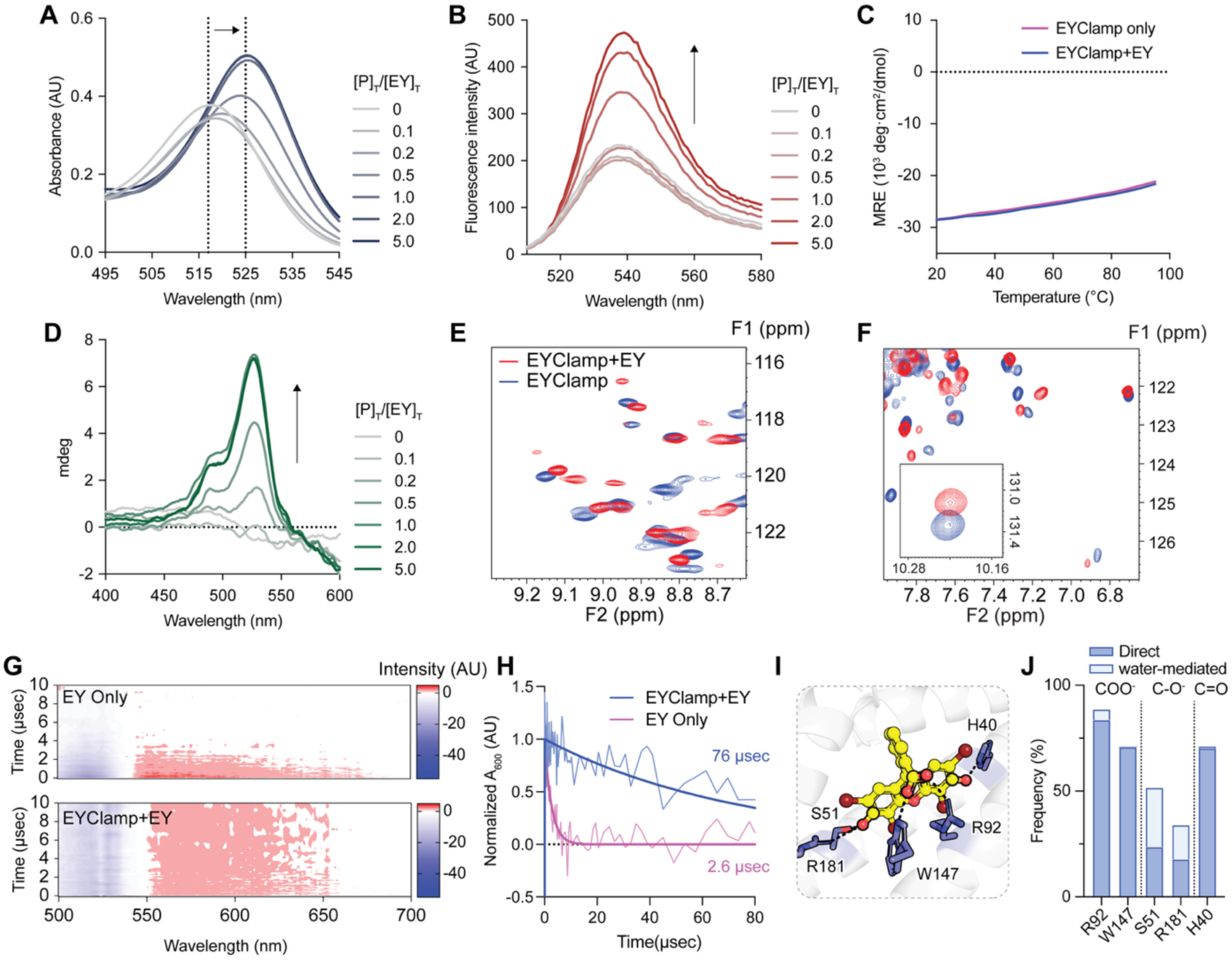
Characterization of fine-tuned EYClamp binding and its biophysical properties. (**A-B**) UV-Vis absorption (**A**) and fluorescence emission (**B**) spectra of EY (1 µM) upon titration with EYClamp, up to a 5-fold molar equivalent of the protein. (**C**) Thermal melting curves monitored via CD analysis in far-UV range with 10 uM of EY with 1 equivalent of EYClamp. The CD signal at 222 nm, reported as mean residue ellipticity (MRE), was recorded in 2°C increments from 20°C to 95°C. (**D**) CD spectra demonstrating the induced chirality of 10 µM EY in the visible region upon titrating EYClamp from 0 to 5 molar equivalents. The signal saturates after addition of a single equivalent of EYClamp. (**E-F**) 2D-^1^H,^15^N-HSQC spectra of EYClamp in the presence (red) and absence (blue) of EY, displaying chemical shift perturbations consistent with EY binding in the fast-exchange regime. The inset provides a magnified view of the tryptophan region. (**G**) Pump-probe nanosecond transient absorption spectra in the UV-Vis regime for free EY (upper) and EY bound to EYClamp (lower). Samples were prepared with either 50 µM free EY or 50 µM EY-EYClamp complex, excited at 530 nm, then probed after varying time delays (shown on the Y-axis). A prominent bleaching of the absorption at 500 – 550 nm was observed, associated with the depletion of the ground-state, along with a new absorption at 500 – 600 nm associated with the triplet (T_1_) excited-state. The ground-state bleaching occurs more rapidly, and the T_1_ state is markedly longer-lived in the EY-EYClamp complex compared to free EY. (**H**) Time-resolved profiles of the normalized transient absorption at 600 nm, capturing the T_1_ excited-state dynamics of the EY. (**I**) Structural visualization of the residues in EYClamp that form hydrogen bonds to EY. (**J**) Fraction of intermolecular hydrogen bonds formed between EYClamp and EY during the last 500 nsec of a 1 μsec unbiased all-atom MD simulation.

To probe further the native-like structure of EYClamp in the presence and absence of EY, we recorded 2D ^1^H,^15^N-heteronuclear single-quantum coherence (HSQC) NMR spectra. The spectra are reasonably well-resolved and dispersed for a 200-residue α-helical protein, and the addition of EY induced significant spectral shifts, as expected for specific binding in a well-defined pocket (**Figure 2E-F** and **Supplementary Figure 2C**). Although we did not obtain sequence-specific assignments, a single Trp residue in the binding pocket (**Figure 2F inset**) exhibited substantial chemical shift perturbations (ΔδHn > 0.2 ppm or ΔδN > 0.4 ppm).

Upon light excitation, EY forms a singlet excited-state, which relaxes on an ∼1 nsec time scale to a triplet-state. We reason that this is the exact triplet-state that acts as a sensitizer for the reactive species formation from photo-probes, followed by protein labeling. Thus, we used transient absorption spectroscopy (TAS) and time-resolved photoluminescence measurements to interrogate the excited-state dynamics of EY in solution and upon binding EYClamp. The lifetime of the T_1_ excited-state is dramatically increased in the bound complex: free EY exhibits a broad T_1_→T_n_ excited-state absorption (ESA) centered at 600 nm (ρ_T1_ = 2.6 µsec); Upon binding to EYClamp, the EY T_1_ state lifetime increases 29-fold (ρ_T1_ = 76 µsec; **Figure 2G-H** and **Supplementary Figure 2D-E**), providing a longer time window to for reaction with probe molecules.

To elucidate the atomic-level interactions within the binding pocket, we performed unbiased all-atom molecular dynamics (MD) simulations. As the initial model for these simulations, we utilized the Chai-1 predicted structure of the EYClamp-EY complex, which exhibited high structural confidence with a predicted template modeling (pTM) score exceeding 0.95 (**Supplementary Figure 2F**). Analysis of the last 500 nsec of the 1 µsec simulation trajectory revealed that the critical binding interactions are highly stable (**Supplementary Figure 2G-I**). Specifically, hydrogen-bond interactions involving R92, W147, and the rationally designed substitution H40 were maintained with over 70% occupancy, which formed a triad that clamps EY into the pocket (**Figure 2I-J**). Additionally, residues S51 and R181 formed water-mediated hydrogen bonds in the EYClamp, while only R181 formed a hydrogen bond to EY in EYB2.3 (**Supplementary Figure 2J**). Taken together, these data confirm that EYClamp provides a highly stable, structured scaffold that specifically encapsulates EY, therefore rigidifying the ligand in a protected microenvironment and drastically prolonging its excited-state lifetime.

### EYClamp triggers selective target labeling in live cells

We next evaluated the capacity of EYClamp to mediate photo-PLP in live cells. We first transfected EYClamp to HEK293T cells, incubated with EY, washed to remove excessive non-binding EY, then irradiated with LED light in the presence of all tested photo-probes, aryl-diazirine-biotin, aryl-azide-biotin and phenol-biotin (**Supplementary Figure 3A**). We found that EYClamp-expressing cells induced significantly more cellular labeling with all three photo-probes compared to cells without EYClamp (**Supplementary Figure 3A**). As a negative control, we also constructed a disruptive R92W mutant showing at least 100-fold weaker binding to than EYClamp, which we named as “dead EYClamp” (dEYClamp) (**Supplementary Figure 3B**). We observed selective co-elution of EY with EYClamp and virtually no signal with dEYClamp by size-exclusion chromatography (**Supplementary Figure 3C**). We further showed that only purified EYClamp with EY bound (holo-EYClamp) activates significant self-labeling in the presence of light and photo-probes (aryl-diazirine-biotin, aryl-azide-biotin, hydrazide-biotin, phenol-biotin) (**Supplementary Figure 3D**). In contrast, dEYClamp or apo-EYClamp yielded negligible self-labeling. This selective labeling was echoed in the experiment where purified EYClamp or dEYClamp was spiked into cell lysates **Supplementary Figure 3E**). EYClamp drove significant amount of labeling upon either blue or green light irradiation aligns better with the bandwidth of EY.^37^

We then designed a panel of constructs fusing EYClamp or dEYClamp to a non-interacting model protein, EGFP, and developed a workflow for testing photocatalytic self-labeling (**Figure 3A** and **Supplementary Figure 4A**). Each of the fusions were transiently expressed in A549 cells at comparable levels with proper EGFP folding as judged by flow cytometry and Western blotting (**Figure 3B, Supplementary Figure 4A**). We then performed photo-PLP and observed a significant amount of biotinylation by Western blotting for all three photo-probes in EYClamp-EGFP expressing cells but not dEYClamp controls (**Supplementary Figure 4B**). We also tested a genetically encoded HaloTag construct we had previously shown in the ecto-tag MultiMap strategy.^38^ The level of biotinylation for EY-EYClamp was comparable to the covalent HaloTag samples conjugated with the chloroalkane EY derivative ligand, showing that covalent attachment was not much more efficient than tight binding of EY to the EYClamp (**Supplementary Figure 4B**).

**Figure 3:**
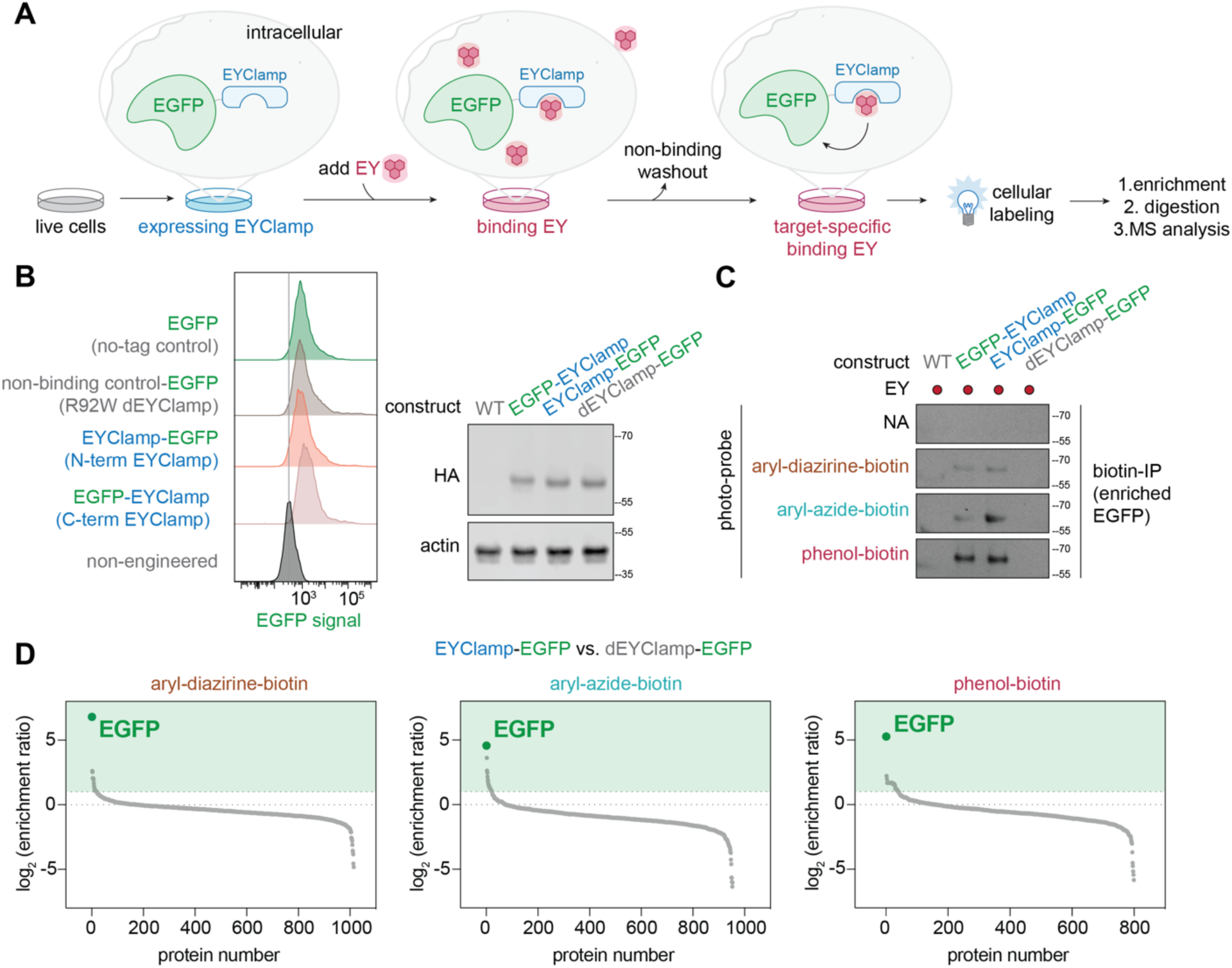
Selective self-labeling of EGFP using EYClamp fusions in live cells. (**A**) Schematic workflow of EYClamp-mediated photo-PLP in live cells. (**B**) Evaluation of expression levels of model protein EGFP and EYClamp fusion constructs using flow cytometry (left) and Western blotting (right). (**C**) Photocatalytic labeling of EGFP using EYClamp fusion designs. Significant enrichment was achieved when EYClamp was fused to either N- or C-terminus but not for dEYClamp. (**D**) Waterfall plot showing EGFP is the top enriched protein for photo-PLP in cells expressing EYClamp-EGFP in comparison to dEYClamp-EGFP in the presence of three photo-probes, aryl-diazirine-biotin, aryl-azide-biotin or phenol-biotin using quantitative proteomics. Significantly enriched proteins are highlighted in the green box with [log_2_(enrichment ratio) **≥** 1]. Data is tabulated in **Supplementary Data Table 2-4.**

We then characterized the selectivity of EYClamp-EGFP self-labeling in cells using photo-PLP with subsequent enrichment (**Figure 3C**). The N-term or C-term EYClamp-EGFP fusions showed comparable photo-induced biotinylation with all three photo-probes, indicating the orientation was not critical. No EGFP was enriched in the dEYClamp-EGFP fusion sample, confirming the requirement of EY binding for target self-labeling. We then performed quantitative proteomics to identify proteome-wide biotinylated proteins using a multiplexed comparison panel (**Supplementary Figure 4C**). We found that EGFP ranked as the highest enriched protein for all three photo-probes when comparing EYClamp-EGFP to dEYClamp-EGFP. We observed selective enrichment with log_2_(enrichment ratios) of 6.81 for aryl-diazirine-biotin, 4.57 for aryl-azide-biotin and 5.27 for phenol-biotin (**Figure 3D, Supplementary Data Table 2-4**). Similar results were obtained when comparing EYClamp-EGFP samples to non-tagged EGFP samples with log_2_(enrichment ratio) of 6.94, 4.05, 4.87 for the three photo-probes, respectively (**Supplementary Figure 4D-E, Supplementary Data Table 5-7**). We also compared EYClamp-EGFP samples to non-engineered cells with log_2_(enrichment ratio) of 7.64, 5.53, 6.11 for the three photo-probes, respectively (**Supplementary Data Table 8-10**). Volcano plots showed that for all comparisons, EGFP was consistently the single most enriched protein with minimal engagement of other endogenous proteins. These studies demonstrated that EYClamp enables robust and selective self-target labeling in cells (**Supplementary Figure 4E** and **5A-C, Supplementary Data Table 2-10**).

### EYClamp enables mapping of E3 ligase neighbors in live cells

We next applied EYClamp-based photo-PLP to study important functional interactomes for E3 ligases. We chose six E3 ligases of interest, including the well-characterized Kelch-like ECH-associated protein 1 (KEAP1), which is known to regulate responses to oxidative stress via proteasomal degradation,^70–72^ along with MDM2, a well-known negative regulator of tumor suppressor p53 through ubiquitin-dependent proteasomal degradation.^73, 74^ We also included four less characterized E3 ligases that are involved in important recent discoveries including: Ankyrin repeat and SOCS box containing 7 (ASB7) that was recently found to associate with oncogenic progression;^75, 76^ mitochondrial RBR-type E3 ligase PARKIN that was associated to early-onset of autosomal neurodegenerative disorder;^77^ transmembrane RING-type E3 ligase RNF43 that regulates the Wnt signaling^78, 79^ and used for extracellular-targeted protein degradation by AbTAC and PROTAB;^80, 81^ STUB1, also known as CHIP that was recently associated to regulate chaperone homeostasis during immune responses and inflammatory diseases.^82, 83^

We first generated fusion constructs of the six E3 ligases with EYClamp (**Supplementary Figure 6A**) and confirmed that all fusion proteins were robustly expressed in HEK293T cells (**Supplementary Figure 6B**). AlphaFold3 predictions of the fusion constructs suggested that the addition of the EYClamp tag would not perturb the predicted native folding of the ligases with the backbone RMSD of EYClamp-ASB7 vs. untagged ASB7 at 1.11Å and EYClamp-STUB1 vs. untagged STUB1 at 1.43Å (**Supplementary Figure 6C**). We then performed cellular labeling using aryl-diazirine-biotin, aryl-azide-biotin and phenol-biotin for all six cell lines and found significant levels of biotinylation (**Figure 4A**). We confirmed that the labeling pattern was similar to those with the EGFP model and that the aryl-azide-biotin generated more labeling than either aryl-diazirine-biotin or phenol-biotin as seen with the EGFP model.

**Figure 4:**
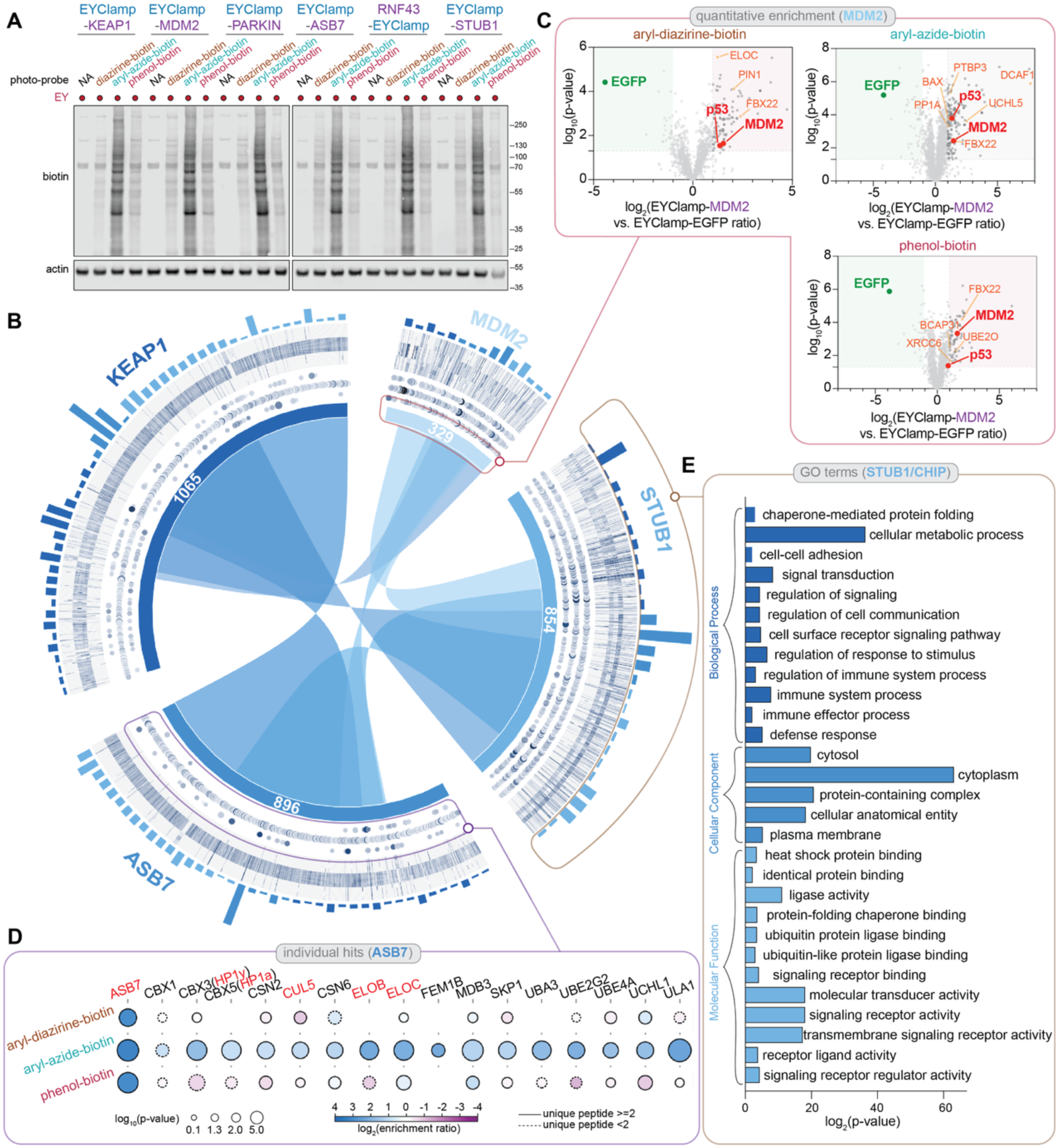
EYClamp is a modular tool that enabled mapping of E3 ligases in live cells. (**A**) E3 ligase fusion construct designs and confirmation of cellular labeling. EYClamp fusion constructs with E3 ligases, KEAP1, MDM2, PARKIN, ASB7, RNF43 and STUB1, were expressed in HEK293T cells and bound to EY. Biotinylation Western blotting analysis showed significant cellular labeling in the presence of photo-probes and light irradiation. (**B**) Chord diagram for E3 ligase interactome analysis using the different EYClamp-E3 ligases. By performing quantitative proteomics of target labeled with EYClamp-KEAP1, EYClamp-MDM2, EYClamp-ASB7 and EYClamp-STUB1, we compared identified neighbors and performed GO analysis. GO terms with high enrichment were displayed including biological processes, cellular component and molecular function (**outer ring**). Data is tabulated in **Supplementary Data Table S29-31** for KEAP1, **Supplementary Data Table S32-34** for ASB7, **Supplementary Data Table S35-37** for MDM2 and **Supplementary Data Table S26-28** for STUB1. Protein-level quantitative enrichment ratios and heatmaps using different probes across four E3 ligases were displayed (**second ring**). Data is tabulated in **Supplementary Data Table S17-19** for KEAP1, **Supplementary Data Table S20-22** for ASB7, **Supplementary Data Table S14-16** for MDM2 and **Supplementary Data Table S23-25** for STUB1. Protein neighborhoods overlap was also displayed (**inner circle**). (**C**) Volcano plots of enriched proteins in cells expressing EYClamp-MDM2 in comparison to EYClamp-EGFP controls using aryl-diazirine-biotin, aryl-azide-biotin, and phenol-biotin, respectively. Significantly enriched proteins are highlighted in red box with [log_2_(enrichment ratio) **≥** 1, P < 0.05, at least 2 unique peptides, three biological replicates]. Data is tabulated in **Supplementary Data Table 14-16**. (**D**) Enrichment profiles of candidate ASB7 neighbors using the three photo-probes. Color (purple to blue) represents increasing log_2_(fold enrichment) whereas size of bubble indicates -log(p-value) across three replicates with larger sizes indicating higher confidence. (**E**) Gene Ontology (GO) analysis performed on all STUB1-interacting candidates. Candidates were compared to UniProt-reviewed human proteome file. Enriched biological process, cellular component and molecular function terms were annotated with -log_10_(p-value). Data is tabulated in **Supplementary Data Table 27-29**.

We then performed biotinylation enrichment and quantitative proteomics upon photo-PLP and analyzed the labeled proteome for four EYClamp-fused E3 ligases: KEAP1, MDM2, STUB1 and ASB7 (**Figure 4B** and **Supplementary Figure 6D-G**). In total, 1633 candidate neighbors were identified across the four E3 ligases providing multitude of information beyond neighborhood identities, including substrate commonality, associated function information and labeling profiles using different probes (**Figure 4B-E**). We first confirmed that the control fusion EYClamp-EGFP preserved its selectivity in EGFP enrichment when expressed in the same plasmid backbone (**Supplementary Figure 6D**). We consistently identified the target E3 ligase as one of the most selectively enriched proteins in the volcano plots compared to the EYClamp-EGFP: Using quantitative proteomics thresholds of [log_2_(fold enrichment) ≥ 1, P < 0.05, at least 2 unique peptides, three biological replicates], MDM2 reached enrichment ratios of 2.90 for aryl-diazirine-biotin, 2.77 for aryl-azide-biotin and 3.05 for phenol-biotin (**Figure 4C, Supplementary Data Table 14-16**); KEAP1 reached enrichment ratios of 3.63 for aryl-diazirine-biotin, 8.22 for aryl-azide-biotin and 3.77 for phenol-biotin (**Supplementary Figure 6E, Supplementary Data Table 17-19**); ASB7 reached higher enrichment ratios of 11.14 for aryl-diazirine-biotin, 13.04 for aryl-azide-biotin and 11.02 for phenol-biotin (**Supplementary Figure 6F, Supplementary Data Table 20-22**); STUB1 also reached higher enrichment ratios of 8.07 for aryl-diazirine-biotin, 10.75 for aryl-azide-biotin and 10.07 for phenol-biotin (**Supplementary Figure 6G-H, Supplementary Data Table 23-25**), respectively. These results confirmed that E3 ligases can be selectively self-labeled and captured using EYClamp-mediated live cell proximity labeling. We then performed in-depth bioinformatics analysis on the enriched proteome that engage with KEAP1, MDM2, STUB1 and ASB7. MDM2 had the narrowest interactome network when compared in parallel to the others (**Figure 4B, Supplementary Data Table 14-16**), which is consistent with the fact that MDM2 has evolved to be specialized to engage mostly with p53 family transcription factors.^84^ In fact, p53 was one of the most highly enriched neighbors of MDM2 using the short-ranged probes aryl-diazirine-biotin and aryl-azide-biotin and moderately with the longer range phenol-biotin (**Figure 4C**). We also found substantial overlap of neighborhoods between E3 ligases: neighborhoods of KEAP1 and ASB7, for example, has a major portion overlapping (**Figure 4B, Supplementary Data Table 17-22**), which maps to their shared roles in responding to oxidative and ER stress.^85, 86^ Consistent with our previous studies,^37, 38^ we found increasing number of hits as we went from aryl-diazirine-biotin ∼ phenol-biotin << aryl-azide-biotin. This suggests that it is useful to include all three photo-probes for a more comprehensive neighborhood map (**Supplementary Figure 7A**).

By comparing the heatmaps of enriched proteins using different photo-probes, we found that aryl-azide-biotin labels more unique targets engaging with KEAP1 and ASB7 than those engaging with STUB1 and MDM2 (**Figure 4B** and **Supplementary Figure 7A, Supplementary Data Table 23-25**). This is possibly due to the larger size of KEAP1 and ASB7, which serve as substrate adapters within large multi-unit Cullin-RING complexes;^75^ on the other hand, MDM2 and STUB1 can function as stand-alone E3 enzymes.^87, 88^ Importantly, ASB7 was only recently found in a CRISPR screen^76^ that it regulates H3K9me3 modification by forming a complex with CUL5. HP1 proteins were found to engage with ASB7 during heterochromatin recruitment using TurboID PLP. When zooming in to the protein identities enriched via EYClamp, we found many of these ASB7-engaging proteins were captured amongst the top candidates, including HP1a (CBX5), HP1y (CBX3), CUL5, Elongin B (ELOB) and Elongin C (ELOC) (**Figure 4D**). Other E3 complex proteins UBA3, UBE4A, UCHL1, ULA1 and the COP9 signalosome proteins CSN2, CSN6 were also found highly enriched. When performing GO analysis, we found biological process terms such as chromatin organization and nuclear transport as well as molecular function terms such as DNA binding and transcription factor activity (**Supplementary Figure 7B, Supplementary Data Table 32-34),** which are closely related to the heterochromatin remodeling functions of ASB7. We then performed GO analysis on KEAP1, MDM2 as well as STUB1, the latter of which was reported with fewer known interactors (**Supplementary Figure 7C-D, Supplementary Data Table 26-31, 35-37**). For STUB1, we found biological process terms such as cell-cell adhesion, immune system processes and signal transduction to be enriched (**Figure 4E**). This observation aligns with recent demonstration of *in vivo* CRISPR screen showing STUB1 as a negative regulator of anti-tumor CD8+ T cell functions in both murine models and in human.^83^ By engaging transmembrane substrates, complexes formed with this E3 ligase were found to be critical for cytokine receptor homeostasis including IL27Ra. Our analysis further pointed to the function of STUB1, where signaling receptor activity and transmembrane signaling receptor activity apart from ligase activity were found to be highly enriched molecular function terms (**Figure 4E**).

## Discussion

We have developed EYClamp, a *de novo* designed proximity labeling enzyme as a genetically encodable scaffold that was engineered to specifically bind the synthetic photocatalyst Eosin Y (EY) for photo-PLP. Designed as a nanomolar-affinity binder, EYClamp is biophysically fine-tuned, highly bioorthogonal, and capable of activating multiple probes with distinct labeling radius. This multi-scale advantage enabled us to monitor protein neighborhoods from a single construct and provide complimentary proximity data at different spatial resolutions and with temporal light control. By showcasing its utility in capturing six important E3 ligases and their neighbors, we successfully mapped their interactomes using a streamlined workflow, which can be harnessed to expand the targetable proteome and accelerate target identification.

From a structural design perspective, a primary advantage of employing a domain-swapped architecture is its potential to generate a highly generalizable scaffold capable of binding diverse co-factors. The resulting inter-domain cavity can be precisely tuned to match the shape complementarity of a given target molecule. In the present work, we used an initially symmetric backbone, which was then diversified into an asymmetric architecture through the addition of specific structural motifs. By modulating the interdomain spacing, it should be possible to accommodate a range of complex and differently sized ligands, as is commonly seen in nature. Furthermore, because these parameterized four-helix bundles inherently support numerous internal binding pockets, the independent diversification of the two swapped domains opens exciting avenues for multi-functional design. This modularity establishes a robust framework for design of proteins with multiple binding sites for novel catalytic and allosteric functions.

We applied EYClamp to E3 ligases given their imperative roles regulating protein homeostasis and cellular functions. With ∼600 E3 ligases encoded by human genome in total, almost all human proteins are regulated by E3 ligases through direct engagement or indirect network regulation via ubiquitin signaling events,^89–93^ all of which steps rely on proximity-induced interactions, including substrate recognition, ubiquitylation, and proteasomal degradation.^66, 94, 95^ While traditional methods such as genetic screening or ubiquitin profiling generate important functional information of these E3 ligases target-by-target,^96–99^ broad interactome profiling has yet to be applied for high-confidence parallel comparison.

We fused the EYClamp to six important E3 ligases, all of which could be robustly expressed in cells and biotinylate proteins using the three photo-probes. We successfully mapped in detail the interactomes using a streamlined workflow, and identified over 1,500 candidate neighbors for KEAP1, MDM2, ASB7 and STUB1. This unbiased mapping revealed critical interactors and highlighted previously uncharacterized functional networks for these targets. Importantly, we identified HP1a (CBX5), HP1y (CBX3), CUL5, ELOB and ELOC along with E3 complex proteins such as UBA3 and UBE4A as ASB7 neighbors using multiple photo-probes with different labeling radius. We also enriched biological processes including chromatin organization, suggesting ASB7 engagement in nuclear regulation. For STUB1, we associated it with cell-cell adhesion and immune system processes, which demonstrated that EYClamp can be harnessed to study functional proteins with distinct localization and activities.

Given the modularity of EYClamp and it being assembled by simply binding off-the-shelf EY, we envision that many previously elusive proximity-induced events can be recorded with the assistance of EYClamp. By integrating this genetically encoded tool with our established spatiotemporal mapping workflow,^38^ one should be allowed to track the destiny of the POI however it travels. It will be particularly helpful uncovering new interactions at organelles of harsh environment or cell-cell synapses^23, 100–104^ or shedding light on the mechanisms behind protein homeostasis and translocation.^39, 105, 106^ On one hand, by incorporating various smart logic gates,^107–109^ EYClamp can be incorporated to study complex biological systems including functional synapses or *in vivo*.^31, 110–113^ On the other hand, similar to successful demonstration of BioTAC,^114^ E-STUB^115^ and Ub-POD^116^, future work can also use EYClamp to monitor target interactome changes upon drug treatment or mutant regulation,^99, 117–119^ where mechanistic pathways of allosteric E3 ligase regulator can be revealed and evaluated.^120, 121^ We envision EYClamp can be easily fused to any POI in different cell types via genetic manipulation, which can be of great assistance for high-throughput target interactome profiling and functional evaluation before any biochemical validation experiments.

## Acknowledgements

We thank Mark J. S. Kelly, Hyunil Jo, Pornparn Kongpracha and Justin McKetney for technical support and helpful discussions. We are grateful to generous support from NIH-1R01CA248323-01 (J.A.W.), NIH-R35GM122451 (J.A.W.), NIH-R35-GM122603 (W.F.D), the Anne Wojcicki Foundation (J.A.W.), the Hind Professorship in Pharmaceutical Sciences (J.A.W.), NSF-CHE-2528384 (W.F.D), NSF-MCB-2306190 (W.F.D), NSF-CHE-2528383 (M.J.T), NSF-ITE-2448848 (M.J.T.), RS-2025-02213459 (Y.H.K), U54CA274502 (N.J.K), RS-2023-00252972 (N.H.K), and The ministry of Trade Industry, and Energy in Korea, under Fostering Global Talents for Innovative Growth Program (P0008746) (N.H.K). Z.L. is a Shanghai Academy of Natural Sciences (SANS) scholar (Early Career).

