## Supplementary Information for "A *de novo* designed enzyme for photo-proximity labeling of E3 ligase neighborhoods in live cells"

neighborhoods in live cells

**Index**

**Supplementary Figure 1**: Strategy for iterative design and testing of EYClamp using a domain-swapped dimer.
**Supplementary Figure 2**: Biophysical characterization of EYClamp.
**Supplementary Figure 3**: EYClamp activates labeling using different photo-probes in vitro.
**Supplementary Figure 4**: EYClamp expresses and labels in live cells.
**Supplementary Figure 5**: EYClamp when fused to EGFP selectively self-labels in cells.
**Supplementary Figure 6**: EYClamp captured E3 ligases in live cells.
**Supplementary Figure 7**: EYClamp mapping provided functional insights into E3 ligases.

**Supplementary Data Table 1**: Protein sequences used in this study
**Supplementary Data Table 2**: Full list of proteins identified by quantitative proteomics enriched in A549 cells expressing EYClamp-EGFP in comparison to dEYClamp controls using aryl-diazirine-biotin.
**Supplementary Data Table 3**: Full list of proteins identified by quantitative proteomics enriched in A549 cells expressing EYClamp-EGFP in comparison to dEYClamp controls using aryl-azide-biotin.
**Supplementary Data Table 4**: Full list of proteins identified by quantitative proteomics enriched in A549 cells expressing EYClamp-EGFP in comparison to dEYClamp controls using phenol-biotin.
**Supplementary Data Table 5**: Full list of proteins identified by quantitative proteomics enriched in A549 cells expressing EYClamp-EGFP in comparison to no-tag EGFP controls using aryl-diazirine-biotin.
**Supplementary Data Table 6**: Full list of proteins identified by quantitative proteomics enriched in A549 cells expressing EYClamp-EGFP in comparison to no-tag EGFP controls using aryl-azide-biotin.
**Supplementary Data Table 7**: Full list of proteins identified by quantitative proteomics enriched in A549 cells expressing EYClamp-EGFP in comparison to no-tag EGFP controls using phenol-biotin.
**Supplementary Data Table 8**: Full list of proteins identified by quantitative proteomics enriched in A549 cells expressing EYClamp-EGFP in comparison to non-engineered controls using aryl-diazirine-biotin.
**Supplementary Data Table 9**: Full list of proteins identified by quantitative proteomics enriched in A549 cells expressing EYClamp-EGFP in comparison to non-engineered controls using aryl-azide-biotin.

**Supplementary Data Table 10**: Full list of proteins identified by quantitative proteomics enriched in A549 cells expressing EYClamp-EGFP in comparison to non-engineered controls using phenol-biotin.
**Supplementary Data Table 11**: Full list of proteins identified by quantitative proteomics enriched in HEK293T cells expressing EYClamp-EGFP in comparison to non-expressing controls using aryl-diazirine-biotin.
**Supplementary Data Table 12**: Full list of proteins identified by quantitative proteomics enriched in HEK293T cells expressing EYClamp-EGFP in comparison to non-expressing controls using aryl-azide-biotin.
**Supplementary Data Table 13**: Full list of proteins identified by quantitative proteomics enriched in HEK293T cells expressing EYClamp-EGFP in comparison to non-expressing controls using phenol-biotin.
**Supplementary Data Table 14**: EYClamp-enriched protein list in MDM2 expressing cells using aryl-diazirine-biotin.
**Supplementary Data Table 15**: EYClamp-enriched protein list in MDM2 expressing cells using aryl-azide-biotin.
**Supplementary Data Table 16**: EYClamp-enriched protein list in MDM2 expressing cells using phenol-biotin.
**Supplementary Data Table 17**: EYClamp-enriched protein list in KEAP1 expressing cells using aryl-diazirine-biotin.
**Supplementary Data Table 18**: EYClamp-enriched protein list in KEAP1 expressing cells using aryl-azide-biotin.
**Supplementary Data Table 19**: EYClamp-enriched protein list in KEAP1 expressing cells using phenol-biotin.
**Supplementary Data Table 20**: EYClamp-enriched protein list in ASB7 expressing cells using aryl-diazirine-biotin.
**Supplementary Data Table 21**: EYClamp-enriched protein list in ASB7 expressing cells using aryl-azide-biotin.
**Supplementary Data Table 22**: EYClamp-enriched protein list in ASB7 expressing cells using phenol-biotin.
**Supplementary Data Table 23**: EYClamp-enriched protein list in STUB1 expressing cells using aryl-diazirine-biotin.
**Supplementary Data Table 24**: EYClamp-enriched protein list in STUB1 expressing cells using aryl-azide-biotin.
**Supplementary Data Table 25**: EYClamp-enriched protein list in STUB1 expressing cells using phenol-biotin.
**Supplementary Data Table 26**: Biological Process Gene Ontology (GO) terms enriched with STUB1-interacting candidates.
**Supplementary Data Table 27**: Cellular component Gene Ontology (GO) terms enriched with STUB1-interacting candidates.
**Supplementary Data Table 28**: Molecular function Gene Ontology (GO) terms enriched with STUB1-interacting candidates.
**Supplementary Data Table 29**: Biological Process Gene Ontology (GO) terms enriched with KEAP1-interacting candidates.
**Supplementary Data Table 30**: Cellular component Gene Ontology (GO) terms enriched with KEAP1-interacting candidates.

**Supplementary Data Table 31**: Molecular function Gene Ontology (GO) terms enriched with KEAP1-interacting candidates.
**Supplementary Data Table 32**: Biological Process Gene Ontology (GO) terms enriched with ASB7-interacting candidates.
**Supplementary Data Table 33**: Cellular component Gene Ontology (GO) terms enriched with ASB7-interacting candidates.
**Supplementary Data Table 34**: Molecular function Gene Ontology (GO) terms enriched with ASB7-interacting candidates.
**Supplementary Data Table 35**: Biological Process Gene Ontology (GO) terms enriched with MDM2-interacting candidates.
**Supplementary Data Table 36**: Cellular component Gene Ontology (GO) terms enriched with MDM2-interacting candidates.
**Supplementary Data Table 37**: Molecular function Gene Ontology (GO) terms enriched with MDM2-interacting candidates.

**Methods:**

General methods and instrumentation.
Antibodies and biological reagents.
General chemical methods and instrumentation.
Overview of design process.
Construct designs.
Protein expression and purification.
HSQC measurement.
Fluorescence emission and polarization measurement for EY binding assessment.
Circular dichroism measurement.
Pump-probe transient UV/Vis spectroscopy.
Unbiased classical molecular dynamics analysis.
Western blot protocol.
Mammalian cell culture and transfection.
EYClamp labeling assay using purified proteins.
EYClamp labeling assay in cells.
Biotinylated protein enrichment.
Sample preparation for LC-MS/MS analysis.
Proteomics analysis of digested peptide samples.
Analysis of proteomics datasets.
Software.
Statistics and Reproducibility.
References.

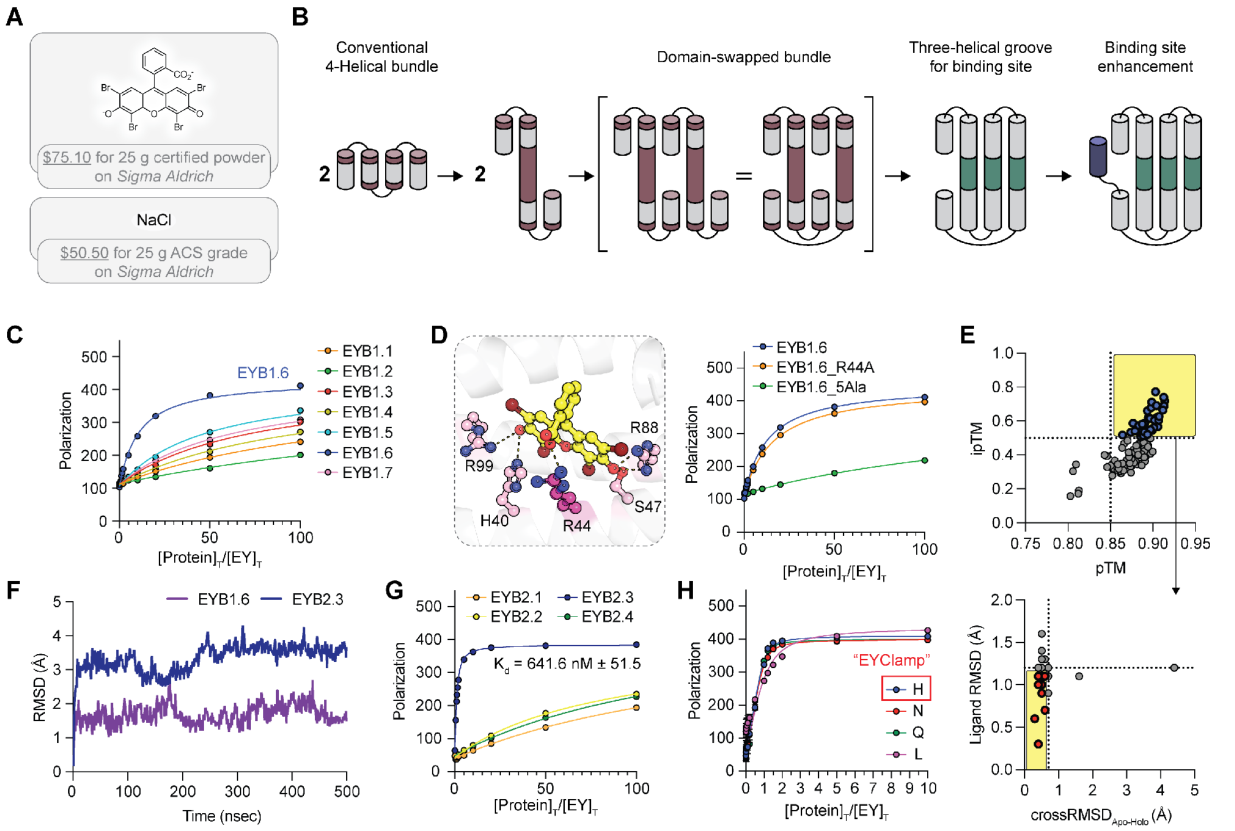


**Supplementary Figure 1: Strategy for iterative design and testing of EYClamp using a domain-swapped dimer. (A)** The Eosin Y (EY) catalyst is as cheap as reagent-grade sodium chloride. **(B)** Schematic representation of the domain-swapped dimerization approach. **(C)** Binding affinities of the EYB1 variants measured via a fluorescence polarization assay. The protein was titrated from 0 to 100 equivalent in the presence of 1 μM of EY. **(D)** Model of specific binding residues interacting with EY in EYB1.6, alongside binding profiles of the EYB1.6 variants. The complex structure was predicted by Chai-1. EYB1.6_R44A denotes a variant with an R44A mutation, while EYB1.6_5Ala represents a variant where all five residues (H40, R44, S47, R88, and R99) expected to interact with EY are mutated to alanine. **(E)** Two-step filtering of 100 sequences generated by LigandMPNN for the 20-amino-acid extended scaffold using Chai-1 predictions. **(upper panel)** In the first step, design candidates were filtered by structural confidence scores, with the x-axis representing the interface predicted template modeling score (ipTM) and the y-axis representing the overall predicted template modeling score (pTM). Sequences passing the criteria of pTM > 0.85 and ipTM > 0.5 were retained. **(lower panel)** In the second step, the remaining candidates were filtered by structural convergence across 10 generated models per sequence. The x-axis represents the ligand root-mean-square deviation (RMSD), and the y-axis represents the cross-state RMSD between the predicted apo- and holo- models. Final design candidates were selected using strict thresholds of a cross-state RMSD < 0.7 Å and a ligand RMSD < 1.2 Å. **(F)** Root-mean-square deviation (RMSD) analysis from unbiased all-atom MD simulations of EYB1.6 and

EYB2.3. **(G)** Binding profiles of the EYB2 variants measured via FP. **(H)** Binding profiles of EYB2.3 variants incorporating diverse hydrogen-bond donor mutations at position 40.

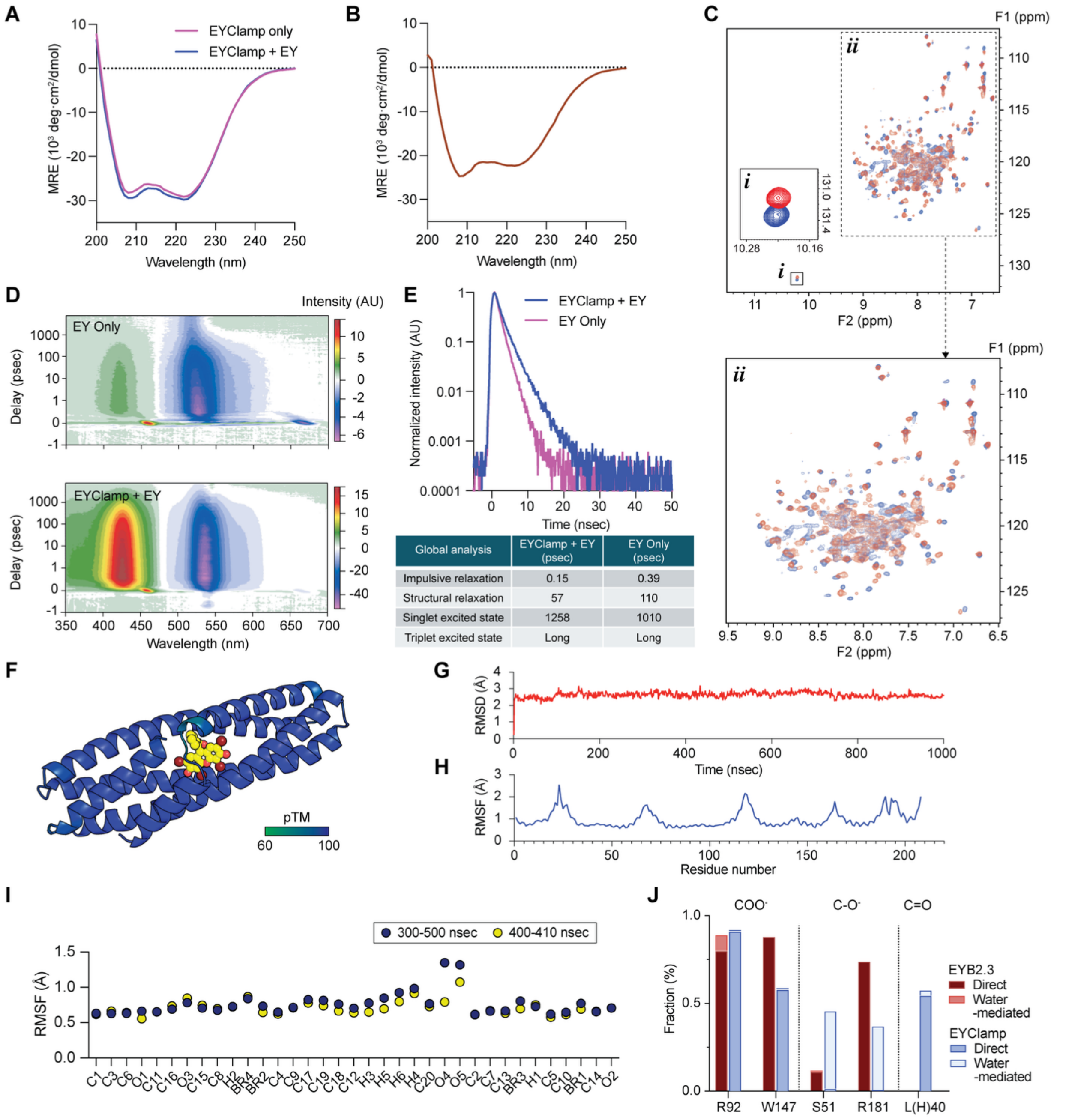


| Global analysis | EYClamp + EY (psec) | EY Only (psec) |
|---|---|---|
| Impulsive relaxation | 0.15 | 0.39 |
| Structural relaxation | 57 | 110 |
| Singlet excited state | 1258 | 1010 |
| Triplet excited state | Long | Long |

**Supplementary Figure 2: Biophysical characterization of EYClamp. (A-B)** Characterization of the helical propensity and thermal stability of EYClamp in the apo- and EY-bound (holo-) states using circular dichroism (CD). **(A)** Near-UV CD spectra of EYClamp in the apo- and holo-states. **(B)** Near-UV CD spectra of EYClamp acquired at 95°C. **(C)** $^1$H-$^{15}$N HSQC spectrum of EYClamp in the absence (blue) and presence (red) of EY. inset box is zoomed in range of tryptophan **(i)** and amide backbone **(ii)**. **(D)** Ultrafast transient absorption spectra in the UV-Vis regime free EY (upper) and for EY when bound to EYClamp (lower). **(E)** Decay profiles measured at the respective emission maxima of EYClamp (Navy) and free EY (purple), capturing the $S_1$ excited-state dynamics. **(F)** Structural visualization of the predicted EYClamp model, colored by per-residue pTM scores (from 60 as green to 100 as blue). **(G–H)** Root-mean-square deviation **(G)** and root-mean-square fluctuation **(H)** Trajectory analyses during 300 – 500 nsec trajectory from the unbiased all-atom MD simulation of EYClamp. **(I)** Root-mean-square fluctuation (RMSF) analysis of EY in complex with EYClamp from the unbiased all-atom MD simulation. The x-axis represents the atom indices of EY. Fluctuations are represented across different time ranges: 300-500 nsec (blue) and 400-410 nsec (yellow). **(J)** Comparison of intermolecular hydrogen bond frequencies between EYB2.3 and EYClamp.

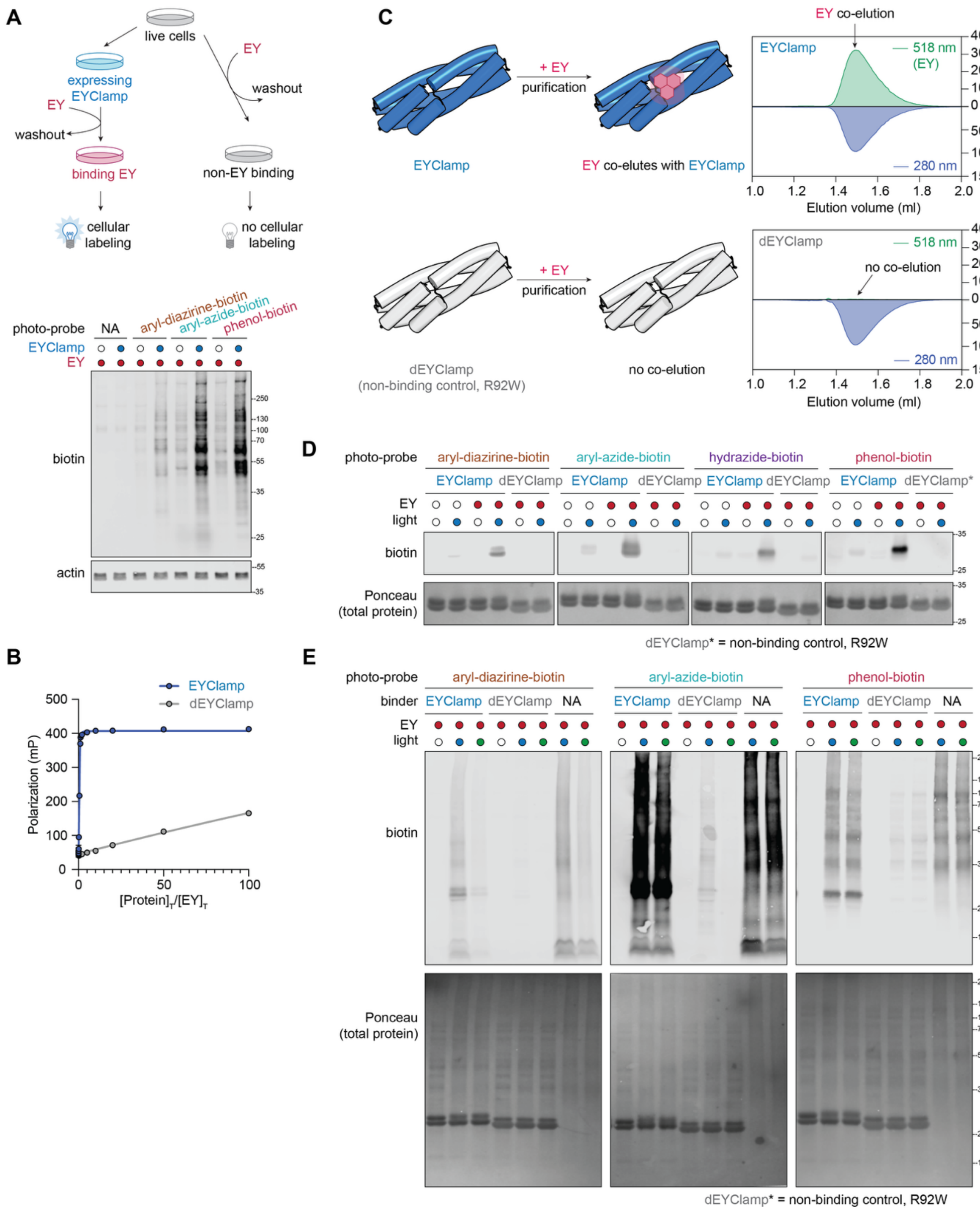


**Supplementary Figure 3: EYClamp activates labeling using different photo-probes in vitro. (A)** Workflow used to test EYClamp cellular labeling using three photo-probes, aryl-diazirine-biotin, aryl-azide-biotin and phenol-biotin. With a one-step EY binding and washing, we can perform photocatalytic protein labeling in cells expressing EYClamp as shown by Western blotting. **(B)** Binding profiles of EYClamp and dEYClamp, determined by titrating the protein (0 to 100 molar equivalents) against a 1 μM EY solution. **(C)** Size-exclusion chromatography (SEC) profiles of EYClamp with EY, which validates complex co-elution. The protein of interest (EYClamp or dEYClamp) and EY were detected via absorbance at 280 nm and 518 nm, respectively. **(D)** Self-biotinylation activated by purified EYClamp. Purified EYClamp with EY bound triggered labeling with aryl-diazirine-biotin, aryl-azide-biotin, hydrazide-biotin and phenol-biotin in the presence of light while dEYClamp and EY-free EYClamp presented minimal background labeling. **(E)** Biotinylation activated by purified EYClamp in cell lysate. Purified EYClamp or dEYClamp was added to cell lysate. Labeling with EYClamp upon blue or green light was observed. Free EY at the same concentration was added as positive controls.

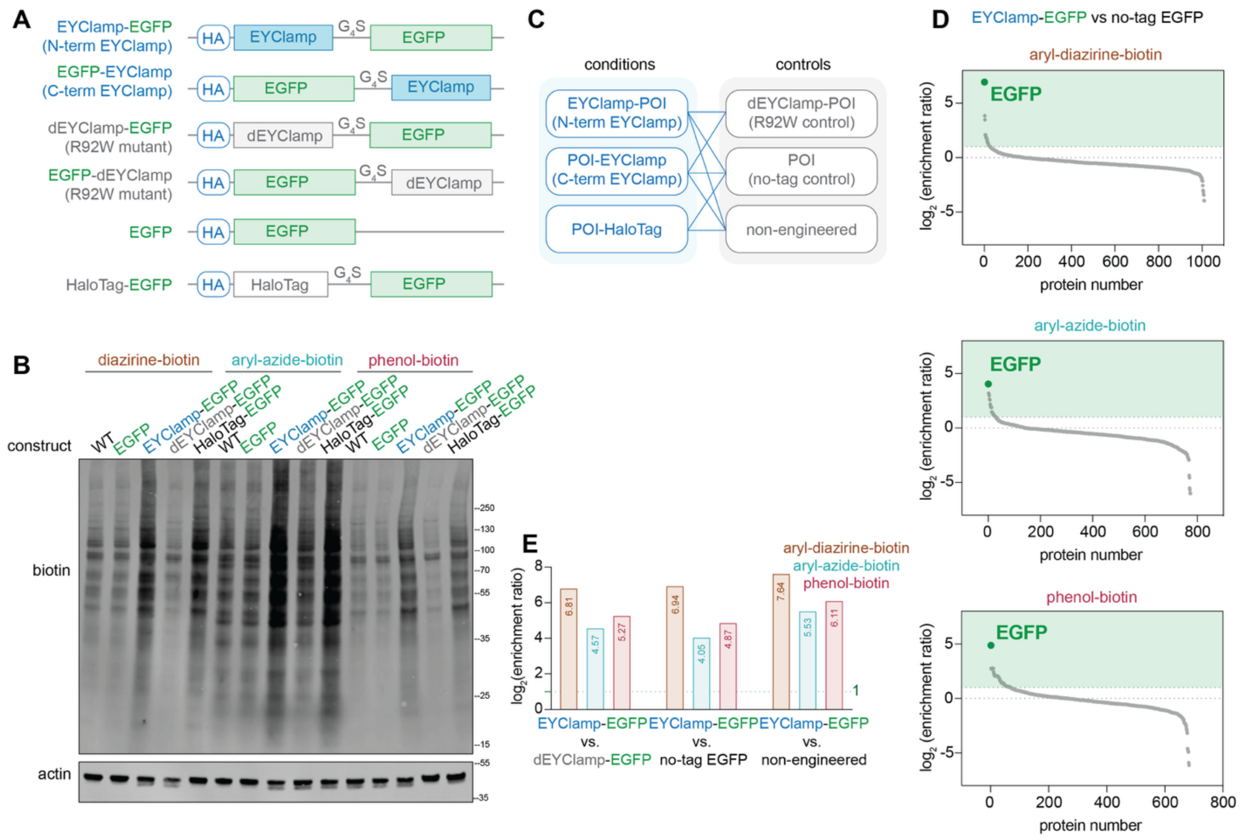


**Supplementary Figure 4: EYClamp expresses and labels in live cells. (A)** Construct designs of EYClamp as well as HaloTag fused to EGFP expressed in A549 and HEK293T cells. **(B)** Western blotting analysis to detect overall biotinylation in A549 cells expressing different constructs in the presence of diazirine-biotin, aryl-azide-biotin and phenol-biotin. **(C)** Full panel of conditions and controls used to study target labeling selectivity. **(D)** Waterfall plot showing EGFP is the top enriched protein for photo-PLP in cells expressing EYClamp-EGFP in comparison to no-tag EGFP control in the presence of three photo-probes, aryl-diazirine-, aryl-azide- or phenol-biotin using quantitative proteomics. Significantly enriched proteins are highlighted in the green box with [$\log_2$(enrichment ratio) ≥ 1] and tabulated in **Supplementary Data Table 5-7**. **(E)** Quantitative enrichment ratios of EGFP in EYClamp-EGFP in comparison to dEYClamp-EGFP, no-tag EGFP and non-engineered controls using aryl-diazirine-biotin, aryl-azide-biotin and phenol-biotin.

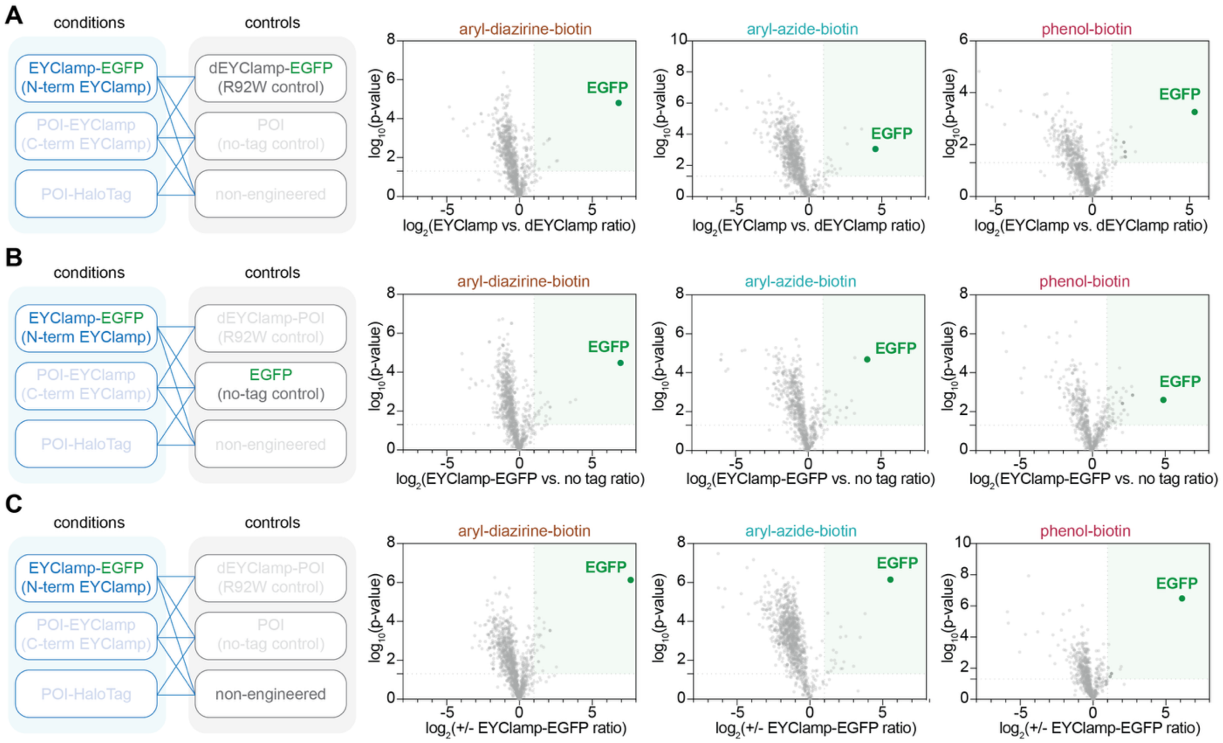


**Supplementary Figure 5: EYClamp when fused to EGFP selectively self-labels in cells. (A)** Volcano plots of enriched proteins in cells expressing EYClamp-EGFP in comparison to dEYClamp-EGFP controls using aryl-diazirine-biotin, aryl-azide-biotin, and phenol-biotin photo-probes, respectively, as shown in **Figure 3D**. Significantly enriched proteins are highlighted in the green box with [$\log_2$(enrichment ratio) ≥ 1, $P < 0.05$, at least 2 unique peptides, three biological replicates]. Data is tabulated in **Supplementary Data Table 2-4**. **(B)** Volcano plots of enriched proteins in cells expressing EYClamp-EGFP in comparison to dEYClamp controls using aryl-diazirine-biotin, aryl-azide-biotin, and phenol-biotin, respectively, as shown in **Supplementary Figure 4D**. Significantly enriched proteins are highlighted in the green box with [$\log_2$(enrichment ratio) ≥ 1, $P < 0.05$, at least 2 unique peptides, three biological replicates]. Data is tabulated in **Supplementary Data Table 5-7**. **(C)** Volcano plots of enriched proteins in cells expressing EYClamp-EGFP in comparison to non-engineered controls using aryl-diazirine-biotin, aryl-azide-biotin, and phenol-biotin, respectively. Significantly enriched proteins are highlighted in the green box with [$\log_2$(enrichment ratio) ≥ 1, $P < 0.05$, at least 2 unique peptides, three biological replicates]. Data is tabulated in **Supplementary Data Table 8-10**.

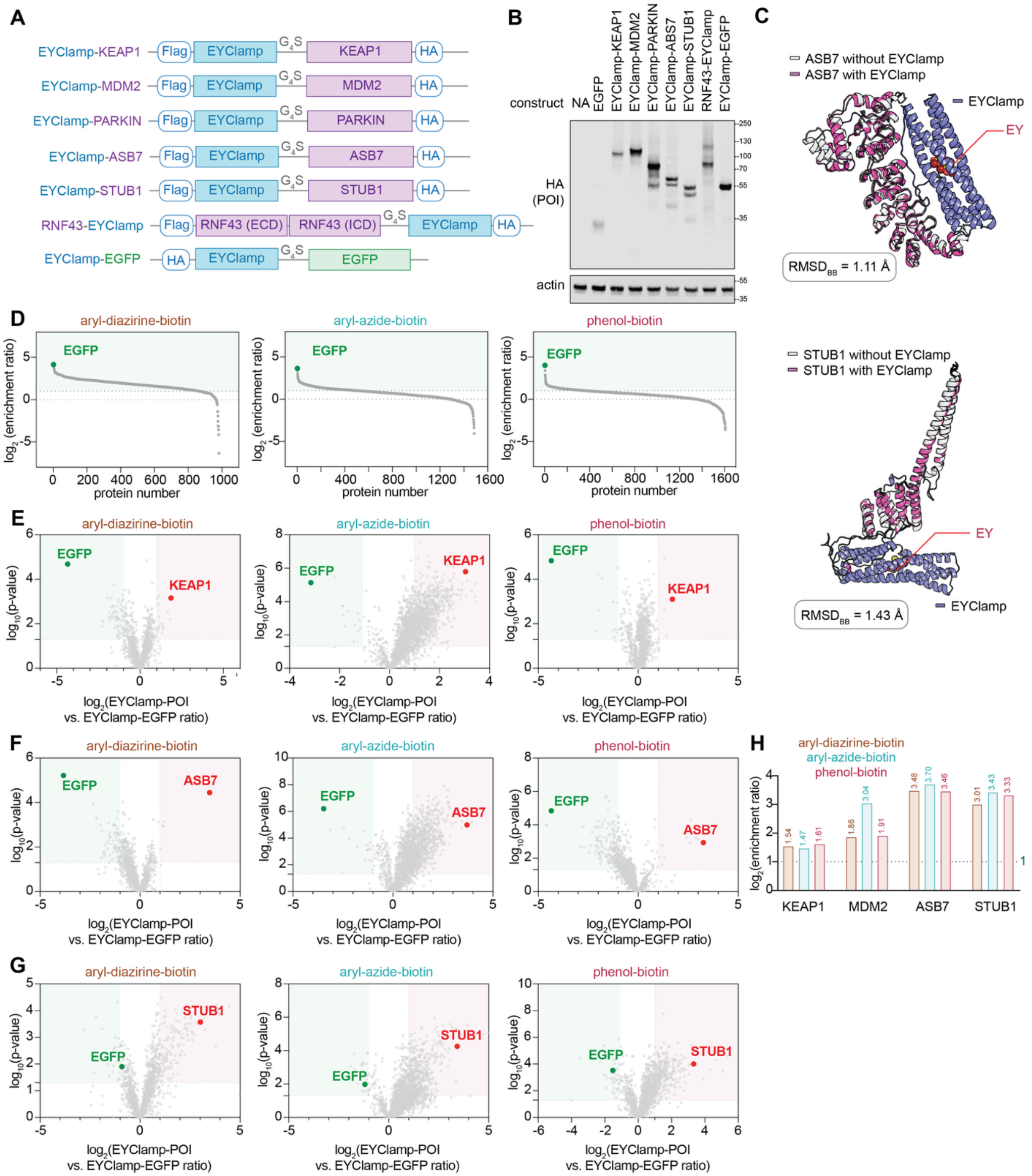


**Supplementary Figure 6: EYClamp captured E3 ligases in live cells. (A)** Construct designs of EYClamp fusion E3 ligase proteins in lentivirus backbones. EYClamp-EGFP was used as a control designed in the same backbone. **(B)** Expression confirmation of E3 ligase-EYClamp fusion proteins in HEK293T cells. **(C)** Structural comparison of E3 ligases and their EYClamp fusions using deep-learning-based prediction models. The E3 ligases alone were predicted using AlphaFold3, whereas the EYClamp-E3 ligase fusion proteins complexed with EY were predicted using Chai-1. Root-mean-square deviation (RMSD) values were calculated to assess structural differences between the isolated E3 ligases and their corresponding EYClamp fusions. **(D)** Ranks of proteins enriched in cells expressing EYClamp-EGFP in comparison to non-engineered control in the presence of aryl-diazirine-biotin, aryl-azide-biotin or phenol-biotin from using quantitative proteomics. EGFP showed up as the most enriched protein throughout conditions. Significantly enriched proteins in EYClamp-EGFP are highlighted in the green box with [$\log_2$(enrichment ratio) ≥ 1, $P < 0.05$, at least 2 unique peptides, three biological replicates]. Data is tabulated in **Supplementary Data Table 11-13**. **(E)** Volcano plots of enriched proteins in cells expressing EYClamp-KEAP1 in comparison to EYClamp-EGFP controls using aryl-diazirine-biotin, aryl-azide-biotin, and phenol-biotin, respectively. Significantly enriched proteins in EYClamp-KEAP1 cells are highlighted in the red box with [$\log_2$ (enrichment ratio) ≥ 1, $P < 0.05$, at least 2 unique peptides, three biological replicates]. Data is tabulated in **Supplementary Data Table 17-19**. **(F)** Volcano plots of enriched proteins in cells expressing EYClamp-ASB7 in comparison to EYClamp-EGFP controls using aryl-diazirine-biotin, aryl-azide-biotin, and phenol-biotin, respectively. Significantly enriched proteins in EYClamp-ASB7 cells are highlighted in the red box with [$\log_2$(enrichment ratio) ≥ 1, $P < 0.05$, at least 2 unique peptides, three biological replicates]. Data is tabulated in **Supplementary Data Table 20-22**. **(G)** Volcano plots of enriched proteins in cells expressing EYClamp-STUB1 in comparison to EYClamp-EGFP controls using aryl-diazirine-biotin, aryl-azide-biotin, and phenol-biotin, respectively. Significantly enriched proteins in EYClamp-STUB1 cells are highlighted in the red box with [$\log_2$(enrichment ratio) ≥ 1, $P < 0.05$, at least 2 unique peptides, three biological replicates]. Data is tabulated in **Supplementary Data Table 23-25**. **(H)** Quantitative enrichment ratios of E3 ligase targets using EYClamp in the presence of aryl-diazirine-biotin, aryl-azide-biotin and phenol-biotin. KEAP1, MDM2, ASB7 and STUB1 were all observed with high enrichment ratios.

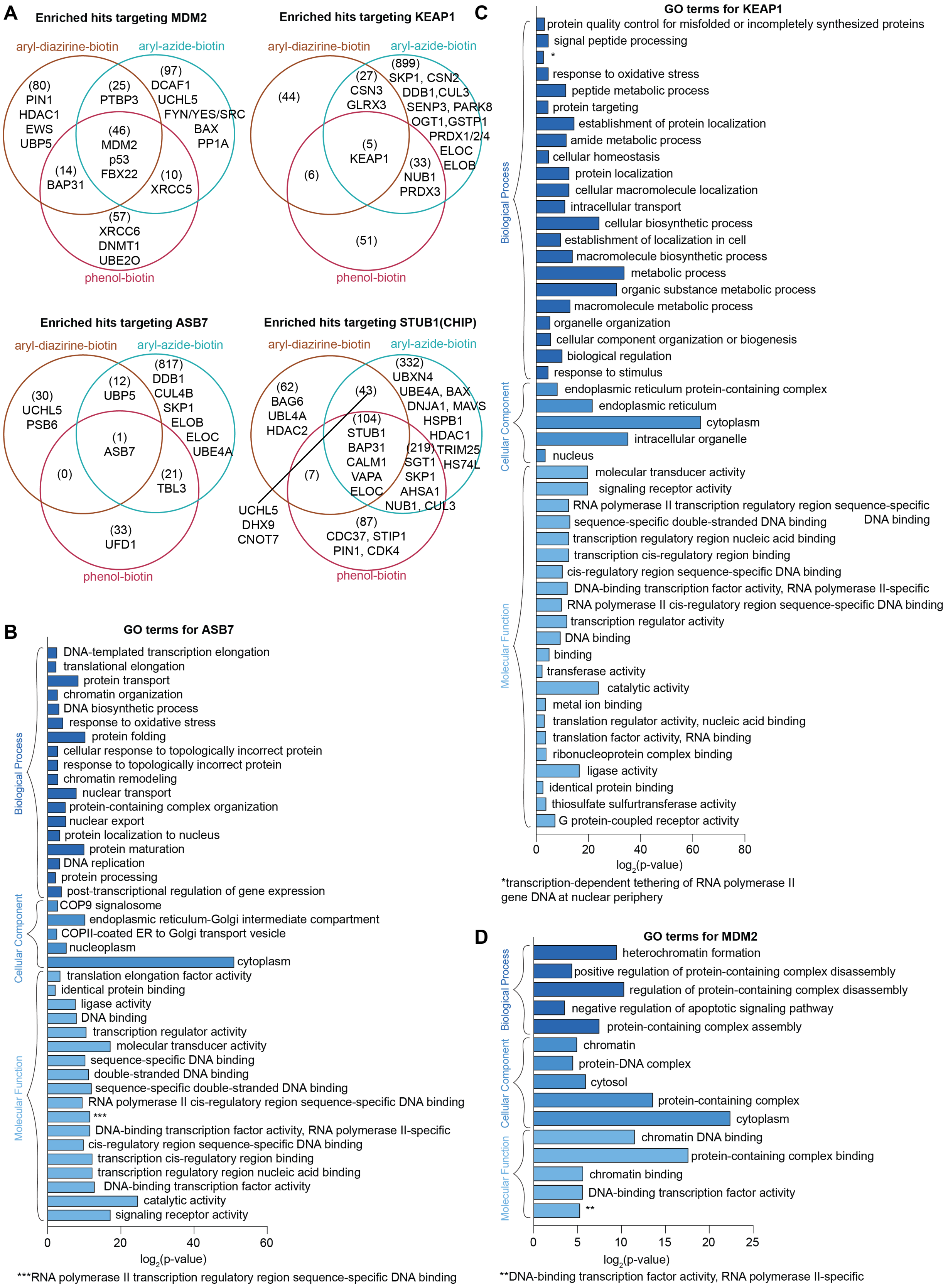
A
Enriched hits targeting MDM2
aryl-diazirine-biotin
aryl-azide-biotin
phenol-biotin
(80) PIN1 HDAC1 EWS UBP5
(25) PTBP3
(97) DCAF1 UCHL5 FYN/YES/SRC BAX PP1A
(46) MDM2 p53 FBX22
(14) BAP31
(10) XRCC5
(57) XRCC6 DNMT1 UBE2O
Enriched hits targeting KEAP1
(44)
(27) CSN3 GLRX3
(899) SKP1, CSN2 DDB1,CUL3 SENP3, PARK8 OGT1,GSTP1 PRDX1/2/4 ELOC ELOB
(5) KEAP1
(6)
(33) NUB1 PRDX3
(51)
Enriched hits targeting ASB7
(30) UCHL5 PSB6
(12) UBP5
(817) DDB1 CUL4B SKP1 ELOB ELOC UBE4A
(1) ASB7
(0)
(21) TBL3
(33) UFD1
Enriched hits targeting STUB1(CHIP)
(62) BAG6 UBL4A HDAC2
(43)
(332) UBXN4 UBE4A, BAX DNJA1, MAVS HSPB1 HDAC1 TRIM25 HS74L
(104) STUB1 BAP31 CALM1 VAPA ELOC
(219) SGT1 SKP1 AHSA1 NUB1, CUL3
(7)
UCHL5 DHX9 CNOT7
(87) CDC37, STIP1 PIN1, CDK4
B
GO terms for ASB7
Biological Process
DNA-templated transcription elongation
translational elongation
protein transport
chromatin organization
DNA biosynthetic process
response to oxidative stress
protein folding
cellular response to topologically incorrect protein
response to topologically incorrect protein
chromatin remodeling
nuclear transport
protein-containing complex organization
nuclear export
protein localization to nucleus
protein maturation
DNA replication
protein processing
post-transcriptional regulation of gene expression
Cellular Component
COP9 signalosome
endoplasmic reticulum-Golgi intermediate compartment
COPII-coated ER to Golgi transport vesicle
nucleoplasm
cytoplasm
Molecular Function
translation elongation factor activity
identical protein binding
ligase activity
DNA binding
transcription regulator activity
molecular transducer activity
sequence-specific DNA binding
double-stranded DNA binding
sequence-specific double-stranded DNA binding
RNA polymerase II cis-regulatory region sequence-specific DNA binding
***
DNA-binding transcription factor activity, RNA polymerase II-specific
cis-regulatory region sequence-specific DNA binding
transcription cis-regulatory region binding
transcription regulatory region nucleic acid binding
DNA-binding transcription factor activity
catalytic activity
signaling receptor activity
0 20 40 60
log2(p-value)
***RNA polymerase II transcription regulatory region sequence-specific DNA binding
C
GO terms for KEAP1
Biological Process
protein quality control for misfolded or incompletely synthesized proteins
signal peptide processing
*
response to oxidative stress
peptide metabolic process
protein targeting
establishment of protein localization
amide metabolic process
cellular homeostasis
protein localization
cellular macromolecule localization
intracellular transport
cellular biosynthetic process
establishment of localization in cell
macromolecule biosynthetic process
metabolic process
organic substance metabolic process
macromolecule metabolic process
organelle organization
cellular component organization or biogenesis
biological regulation
response to stimulus
Cellular Component
endoplasmic reticulum protein-containing complex
endoplasmic reticulum
cytoplasm
intracellular organelle
nucleus
Molecular Function
molecular transducer activity
signaling receptor activity
RNA polymerase II transcription regulatory region sequence-specific DNA binding
sequence-specific double-stranded DNA binding
transcription regulatory region nucleic acid binding
transcription cis-regulatory region binding
cis-regulatory region sequence-specific DNA binding
DNA-binding transcription factor activity, RNA polymerase II-specific
RNA polymerase II cis-regulatory region sequence-specific DNA binding
transcription regulator activity
DNA binding
binding
transferase activity
catalytic activity
metal ion binding
translation regulator activity, nucleic acid binding
translation factor activity, RNA binding
ribonucleoprotein complex binding
ligase activity
identical protein binding
thiosulfate sulfurtransferase activity
G protein-coupled receptor activity
0 20 40 60 80
log2(p-value)
*transcription-dependent tethering of RNA polymerase II gene DNA at nuclear periphery
D
GO terms for MDM2
Biological Process
heterochromatin formation
positive regulation of protein-containing complex disassembly
regulation of protein-containing complex disassembly
negative regulation of apoptotic signaling pathway
protein-containing complex assembly
Cellular Component
chromatin
protein-DNA complex
cytosol
protein-containing complex
cytoplasm
Molecular Function
chromatin DNA binding
protein-containing complex binding
chromatin binding
DNA-binding transcription factor activity
**
0 5 10 15 20 25
log2(p-value)
**DNA-binding transcription factor activity, RNA polymerase II-specific

**Supplementary Figure 7: EYClamp mapping provided functional insights into E3 ligases. (A)** Venn diagram of enriched E3 ligase neighborhoods via EYClamp using different photo-probes. **(B)** Gene Ontology (GO) analysis performed on all KEAP1-interacting candidates. Candidates were compared to UniProt-reviewed human proteome file (downloaded from UniProt database). Enriched biological process, cellular component and molecular function terms were annotated with $-\log_{10}$(p-value) and tabulated in **Supplementary Data Table 29-31**. **(C)** Gene Ontology (GO) analysis performed on all ASB7-interacting candidates. Candidates were compared to UniProt-reviewed human proteome file (downloaded from UniProt database). Enriched biological process, cellular component and molecular function terms were annotated with $-\log_{10}$(p-value) and tabulated in **Supplementary Data Table 32-34**. **(D)** Gene Ontology (GO) analysis performed on all KEAP1-interacting candidates. Candidates were compared to UniProt-reviewed human proteome file (downloaded from UniProt database). Enriched biological process, cellular component and molecular function terms were annotated with $-\log_{10}$(p-value) and tabulated in **Supplementary Data Table 35-37**.

**General methods and instrumentation.**
Sonication of cells or protein pellets was performed using a QSonica Sonicator (Q500, QSonica Sonicators), a Scientz Sonicator (JY92-IIN, Scientz) or a Sonic Dismembrator (Model 500, Fisher Scientific). DNA or protein concentrations were measured using a NanoDrop 2000 spectrophotometer (Thermo Scientific) or a NanoOne spectrophotometer (Yooning). Additional protein quantification was performed by bicinchoninic acid assay on a multimode microplate reader (F Plex Infinite 200 PRO or M Plex Infinite 200 PRO, Tecan Trading AG, Switzerland). Fluorescence emission and fluorescence polarization spectra were recorded using a BioTek Synergy Neo2 plate reader. Immunoblots images were captured by an infrared imager (LI-COR Odyssey CLx or BioRad ChemiDoc MP).

For size exclusion chromatography, analytical and preparative gel filtration profiles were obtained using an AKTA FPLC system (GE Healthcare). For preparative protein separation, concentrated samples were injected onto a Superdex 75 Increase 10/300 GL column and eluted with a PBS mobile phase at a flow rate of 0.5 mL/min. UV absorbance was monitored at 280 nm for protein detection and at 518 nm for EY detection. For analytical runs, samples were injected onto a Superdex 75 Increase 5/150 GL column and eluted with PBS at a flow rate of 0.2 mL/min. Illumination was performed as previously described[1,2] using a Penn PhD Photoreactor M2 (Sigma Aldrich, Z744035) with a 450 nm blue light source module (Sigma Aldrich, Z744033) at 100% intensity, or LED array light sources (Thor Labs, LIU470A for 470 nm LED array, LIU525B for 525 nm LED array) along with a LED mounting adapter (AD38).

Proteomics samples were prepared using part of the Preomics iST kit (Preomics iST 96x, P.O.00027) or the EasyPep kit (micro EasyPep MS sample prep kit, Thermo Scientific). Samples were stored in vials (Thermo Scientific SUN-SRi™ Crimp/Snap Cap Vials, 05-704-225) with caps (Millipore Sigma snap ring vial closures, SU860093). All peptides were collected using protein low-bind Eppendorf tubes (1.5 mL Protein LoBind Tubes, Eppendorf, 022431081; 2 mL Protein LoBind Tubes, Eppendorf, 022431102). Before injection, peptides were dried under vacuum using a Genevac EZ-2 centrifugal evaporator or an Eppendorf Concentrator plus. Peptide concentrations were measured using a NanoDrop 2000 spectrophotometer (Thermo Scientific) and further quantified using a Pierce Quantitative Fluorometric Peptide Assay kit (Thermo Scientific, 23290). Proteomics experiments were performed on a TimsTOF PRO (Bruker) equipped with a CaptiveSpray source and a nanoElute System. The peptides were separated on a 25 cm, ReproSil c18 1.5 µM 100 Å column (PepSep, PSC- 25-150-15-UHP-nc).

**Antibodies and biological reagents.**
Antibodies were purchased including: HA (Cell Signaling Technology, 3724), GFP (Cell Signaling Technology, 2037, 2956), β-actin (Santa Cruz Biotechnology, sc-47778; Cell Signaling Technology, 3700), streptavidin-IR800 (Li-COR, Thermo Scientific 21851), streptavidin-HRP (Thermo Scientific, N100), anti-rabbit IR680 secondary antibody (Rockland, 611-144-002), anti-rabbit IR680 secondary antibody (Thermo Scientific, 35569), anti-mouse IR800 secondary antibody (Rockland, 610-145-211) or anti-mouse IR800 secondary antibody (Thermo Scientific, SA5-10036). All primary antibodies were

diluted 1:1000 and secondary antibodies were diluted 1:5000 for Western blot unless otherwise noted.

For protein enrichment, Ni-NTA agarose resin (HisPur, Thermo Scientific, 88221) and Amicon® 10 kDa MWCO centrifugal filters (Millipore Sigma, UFC901008) were purchased. For enrichment assays, Pierce™ Streptavidin Agarose beads (Pierce, 20349) and mini Bio-Spin columns (Bio-Rad, 732-6207) were used. Cell lysis buffer was prepared by diluting from 10X cell lysis buffer (Cell Signaling Technology, 9803S) or from 10X RIPA buffer (EMD Millipore, 20-188) and supplemented with protease inhibitor (Protease Inhibitor Cocktail 100X, Cell Signaling Technology 5871; Roche cOmplete Protease Inhibitor Cocktail, 11697498001; Halt™ Protease Inhibitor Cocktail 100X, Thermo Scientific, 87786). Sample loading buffer was diluted from 4X Laemmli sample loading buffer (Bio-Rad, 161-0747) or 4X LDS sample buffer (Thermo Scientific, NP0007). Ponceau was purchased (Thermo Scientific, A40000279; Beyotime, P0022). For chemiluminescence, SuperSignal™ West Pico PLUS kit (Thermo Scientific, 34580) was used.

**General chemical methods and instrumentation.**
Chemicals were purchased including TFPA-$PEG_3$-biotin (Thermo Scientific, 21303), biotinyl tyramide (Sigma-Aldrich, SML2135; MCE, HY-125658) and Eosin Y (Sigma-Aldrich, E4009; Thermo Scientific, 409430250). Aryl-diazirine-biotin (diazirine-$PEG_3$-biotin) and Eosin Y-HaloTag ligand were synthesized by Medicilon as previously reported.[2] All solvents and reagents were purchased from chemical suppliers (Sigma Aldrich; Acros Organics; Thermo Scientific; VWR Chemicals BDH®) and were used as received unless otherwise noted.

**Overview of design process.**
A library of a small set of antiparallel 4-helix bundles using Crick parameters as previously generated[3] was used as starting scaffolds for generating domain-swapped dimer. Specifically, we sampled parameters on a grid that varied the bundle radius from 7.9 Å to 8.2 Å, and covaried the superhelical phases of two helices by 14°, resulting in bundles with wide interfaces that varied between 108 and 120° (interhelical Cα-Cα distances of ~ 8.2 - 9.8 Å). Two bundles with same parameter are aligned in same principal axis and extended to be overlapped in three central helices.

The three polar chemical groups of EY were considered for binder design: a carboxyl, a hydroxyl, and a ketone. conformations of EY were computed using molecular mechanics (Maestro, Schrödinger), and the lowest-energy structure was selected for design. The carboxyl interaction was constrained to an arginine residue, exploiting the well-characterized bidentate interaction between the arginine guanidinium group and the two carboxylate oxygens. For the hydroxyl and ketone groups, the scaffold library was screened to position suitable polar residues. Only inward-facing residues were considered, identified using the alpha-hull algorithm in COMBS, which defines the protein surface solely from backbone coordinates.

Backbones that yielded satisfactory solutions were selected based on a root-mean-square deviation (RMSD) of less than 1.0 Å between the van der Mers (vdMer) chemical groups and the ligand with . Poses exhibiting no steric clashes between any portion of the drug molecule and the protein backbone were exclusively advanced and ranked by their vdM scores.[3] Loops connecting the helices were introduced leveraging output from the MASTER program.[4] Aside from the core interacting residues derived from the top-scoring vdM poses, the remainder of the sequence was designed using LigandMPNN.[5] From 100 generated sequences, the 7 candidates with the lowest Rosetta energy units (using the Ref2015 energy function) were selected. These candidates were visually inspected to ensure all hydrogen bonds to the ligand's polar groups were fully satisfied.

To further optimize the initial EYA1.6 scaffold, 20-amino-acid structural extensions were generated using the all-atom RFdiffusion model.[6] From a pool of 100 generated scaffolds featuring an extension that crosses to the opposite side of the helical motif, centroid clustering was used to select a representative candidate for downstream sequence design. The EY ligand was then reoriented to direct its carboxyl group toward the protein interior, and the optimal placement for an arginine residue to interact with this internalized carboxyl group was identified using inverse-COMBS.[7] All residues within 6 Å of EY, along with the newly extended helical motif, were redesigned using LigandMPNN. In total, 100 sequences were generated, and their structural feasibility was assessed using Chai-1.[8] For each candidate, 10 models were predicted and filtered based on a predicted TM-score (pTM) > 0.85 and ipTM score > 0.5 and then filtered further with cross-backbone root-mean-square deviation (RMSD) between apo and holo state < 0.7 Å across the 10 models, and a ligand RMSD < 1.2 Å. Final candidates were selected for experimental validation.

To further refine the hydrogen-bonding network of EYB2.3 at residue 40, 1,000 sequence variants were generated using LigandMPNN across variable temperature sampling parameters. The resulting amino acid frequencies were then quantified to establish a residue probability distribution for that position.

**Construct designs.**

For protein expression in *Escherichia coli*, codon-optimized plasmids encoding the designed candidate proteins, featuring an N-terminal 6xHis tag and a TEV protease cleavage site (HHHHHHENLYFQS) were synthesized by Twist Bioscience. Overhang sequences were appended to the 5' (CTCTAGAAATAATTTTGTTTAACTTTAAGAAGGAGATATACC) and 3' (GATCCGGCTGCTAACAAAGCCCGAAAG) ends to facilitate Gibson assembly into the pET-28a (+) vector.

For protein expression in mammalian cells, codon-optimized plasmids encoding EYClamp and control designs were constructed in either a pcDNA3.1(+) backbone or a pGenLenti backbone by standard molecular biology methods as previously reported[9] with GGGGS links in between, which were synthesized by GenScript or General Biosystems. All plasmids were verified by Sanger (Quintarabio or Sangon) or whole-plasmid (Primordium Labs) sequencing.

Plasmids used include: (Plasmid name – backbone)

| | Plasmid name | Backbone |
|---|---|---|
| 1. | pET-28a (+)_His-TEV-EYB1.1 | pET-28a (+) |
| 2. | pET-28a (+)_His-TEV-EYB1.2 | pET-28a (+) |
| 3. | pET-28a (+)_His-TEV-EYB1.3 | pET-28a (+) |
| 4. | pET-28a (+)_His-TEV-EYB1.4 | pET-28a (+) |
| 5. | pET-28a (+)_His-TEV-EYB1.5 | pET-28a (+) |
| 6. | pET-28a (+)_His-TEV-EYB1.6 | pET-28a (+) |
| 7. | pET-28a (+)_His-TEV-EYB1.7 | pET-28a (+) |
| 8. | pET-28a (+)_His-TEV-EYB1.6_R44A | pET-28a (+) |
| 9. | pET-28a (+)_His-TEV-EYB1.6_5Ala | pET-28a (+) |
| 10. | pET-28a (+)_His-TEV-EYB2.1 | pET-28a (+) |
| 11. | pET-28a (+)_His-TEV-EYB2.2 | pET-28a (+) |
| 12. | pET-28a (+)_His-TEV-EYB2.3 | pET-28a (+) |
| 13. | pET-28a (+)_His-TEV-EYB2.4 | pET-28a (+) |
| 14. | pET-28a (+)_His-TEV-EYB2.3_L40Q | pET-28a (+) |
| 15. | pET-28a (+)_His-TEV-EYB2.3_L40N | pET-28a (+) |
| 16. | pET-28a (+)_His-TEV-EYClamp | pET-28a (+) |
| 17. | pET-28a (+)_His-TEV-dEYClamp | pET-28a (+) |
| 18. | pcDNA3.1(+)_HA-EYClamp-EGFP | pcDNA3.1(+) |
| 19. | pcDNA3.1(+)_HA-EGFP-EYClamp | pcDNA3.1(+) |
| 20. | pcDNA3.1(+)_HA-dEYClamp-EGFP | pcDNA3.1(+) |
| 21. | pcDNA3.1(+)_HA-EGFP-dEYClamp | pcDNA3.1(+) |
| 22. | pcDNA3.1(+)_HA-EGFP | pcDNA3.1(+) |
| 23. | pcDNA3.1(+)_HaloTag_EGFP | pcDNA3.1(+) |
| 24. | pcDNA3.1(+)_HA-EYClamp-KEAP1 | pcDNA3.1(+) |
| 25. | pcDNA3.1(+)_HA-EYClamp-ASB7 | pcDNA3.1(+) |
| 26. | pGenLenti_HA-EYClamp-EGFP | pGenLenti |
| 27. | pGenLenti_Flag-EYClamp-KEAP1-HA | pGenLenti |
| 28. | pGenLenti_Flag-EYClamp-MDM2-HA | pGenLenti |
| 29. | pGenLenti_Flag-EYClamp-KEAP1-HA | pGenLenti |
| 30. | pGenLenti_Flag-EYClamp-PARKIN-HA | pGenLenti |
| 31. | pGenLenti_Flag-EYClamp-ASB7-HA | pGenLenti |
| 32. | pGenLenti_Flag-RNF43-EYClamp-HA | pGenLenti |
| 33. | pGenLenti_Flag-EYClamp-STUB1-HA | pGenLenti |

**Protein expression and purification.**

The recombinant pET-28a (+) plasmids were transformed into *Escherichia coli* BL21(DE3) cells. Single colonies from LB agar plates were inoculated into Terrific Broth (TB) supplemented with 50 μg/mL kanamycin and grown overnight as starter cultures. These were transferred into 200 mL of TB medium with kanamycin and grown at 37 °C until reaching an $OD_{600}$ of 0.8. Protein expression was induced with 0.5 mM isopropyl β-D-1-thiogalactopyranoside (IPTG). Following induction, cultures were grown overnight at 30 °C, and cells were harvested by centrifugation.

Cell pellets were resuspended in 25 mL of PBS buffer (10 mM $Na_2HPO_4$, 1.8 mM $KH_2PO_4$, 2.7 mM KCl, 137 mM NaCl, pH 7.4) containing 20 mM imidazole, and lysed via sonication (Sonic Dismembrator Model 500, Fisher Scientific). The lysate was clarified by centrifugation, and the supernatant was applied to a gravity column packed with 1.0 mL or 3.0 mL of Ni-NTA agarose resin (HisPur, Thermo Scientific, 88221). The resin was washed with 3 column volumes (CVs) of PBS buffer containing 20 mM imidazole, and the bound protein was eluted with 7 mL of PBS containing 250 mM imidazole. The eluted proteins were concentrated and buffer-exchanged three times into standard PBS using an Amicon® 10 kDa MWCO centrifugal filter (Millipore Sigma, UFC901008) to remove residual imidazole.

**HSQC measurement.**
Isotopically $^{15}$N-labeled proteins were expressed by inoculating *E. coli* colonies into 20 mL of LB media grown overnight at 37 °C. The starter cultures were diluted 1:50 into 1 L of fresh [U-$^{15}$N]-labeled M9 minimal medium and grown at 37 °C until the $OD_{600}$ reached 0.6–0.8. Expression was induced with a final concentration of 500 μM IPTG, and cultures were grown overnight at 30 °C. The expressed proteins were initially purified using the Ni-NTA pull-down protocol described above, followed by size-exclusion chromatography utilizing an AKTA FPLC system.

NMR samples were prepared at a final protein concentration of 400 μM, supplemented with 5% (v/v) to provide a lock signal. All NMR spectra were acquired at 298.1 K. Two-dimensional (2D) $^1$H,$^{15}$N- HSQC (pulse program: fhsqcf3gpph) spectra were recorded on a Bruker NEO 800 MHz spectrometer with a 5-mm TCI H&F-C/N-D CryoProbe. Spectra were processed using TopSpin 3.6.3 (Bruker).

**Fluorescence emission and polarization measurement for EY binding assessment.**
To assess binding, EY dissolved in deionized water was mixed with purified proteins in PBS buffer and incubated for 5 minutes at room temperature prior to measurement. Fluorescence emission spectra were recorded in black, flat-bottom 96-well plates using a BioTek Synergy Neo2 plate reader with an excitation wavelength of 460 nm. Protein aliquots from 10 or 100 μM stocks in PBS were diluted to formulate 200 μL samples containing 1.0, 0.5, or 0.25 μM EY. Each condition was measured in triplicate. Fluorescence polarization (FP) assays were performed on the same samples using a BioTek Synergy 2 plate reader equipped with 460 nm excitation and 516 nm emission filters. FP values were recorded in polarization (P) units. Polarization values were plotted against protein concentration, and the dissociation constant (*K*d) was determined by fitting the data with the equation below.

$$Y = Y_0 + (Y_{max} - Y_0) \cdot \left(\frac{-b - \sqrt{b^2 - 4 \cdot a \cdot c}}{2 \cdot a \cdot P}\right)$$

Where, a = 1, b = $-K_d$ – P/n – X, c = (P·X)/n (P = total ligand concentration, $K_d$ = dissociation constant, $Y_0$ = signal of the free ligand, Y = observed signal of fluorescence polarity, X = total protein concentration, n = stoichiometry).

**Circular dichroism measurement.**

CD spectra were acquired on a spectropolarimeter (Jasco, J-810). Wavelength scans were performed from 200 to 250 nm to assess protein secondary structure, and from 400 to 600 nm to monitor the induced chirality of Eosin-Y. Protein samples were prepared at 10 μM in PBS buffer and analyzed in a 0.1 cm path-length quartz cuvette. To determine the melting temperature ($T_m$), thermal denaturation was monitored from 20 to 95 °C, utilizing a heating rate of 2 °C/min and recording measurements at 5 °C intervals.

**Pump-probe transient UV/Vis spectroscopy.**
Transient absorption spectra were obtained through standard pump-probe method. Optical pulses (≥ 80 fs, 1 kHz) centered at 800 nm were generated with Solstice Ace (Spectra-Physics, Milpitas, CA, USA), which consisted of a regenerative amplifier seeded by a mode-locked Ti:Sapphire oscillator. A fraction of the output from the regenerative amplifier was split to feed an optical parametric amplifier TOPAS-C (Light Conversion, Vilnius, Lithuania), which generates pump pulses whose center wavelength was tuned for each measurement. The pump beam was fed into HARPIA spectroscopy system (Light Conversion, Vilnius, Lithuania). HARPIA chopper was set at f/4 (250 Hz). Another fraction of the regenerative amplifier output was fed into HARPIA, passed through an 8-ns optical delay line, focused onto a 3 mm calcium fluoride window for supercontinuum generation, and used as a probe beam for ultrafast transient absorption spectroscopy. The probe pulse for nanosecond transient absorption spectroscopy was generated by the fiber laser SuperK COMPACT (NKT Photonics, Birkerød, Denmark). The polarization and attenuation of the pump and probe beams were controlled by half-wave plate and Glan-Taylor prism pairs. Between pump and probe beams, the polarization was set for the magic angle (54.7°). The pump beam was focused onto the sample cuvette with an f = +100 cm lens, while the probe beam was focused with a concave mirror. The spot size diameter was ~0.2 – 0.3 mm. The pump power utilized was about 600 μW for ultrafast transient absorption spectra, and 3000 μW for nanosecond transient absorption spectra. The probe beam after passing through the sample was adjusted with an ND filter to avoid detector saturation. Then the probe beam was focused onto the entrance slit of spectrograph Kymera 328i (Andor Technology, Belfast, UK), which was interfaced by a broadband Si n-channel metal-oxide semiconductor (NMOS) detector (S3901 + C7884, Hamamatsu Photonics, Hamamatsu City, Shizuoka, Japan). All measurements utilized a 2 mm path length quartz sample cell without degassing. All transient spectra reported represent averages obtained over 9 scans, with each scan consisting of over 150 time-delays. Following all pump-probe transient absorption experiments, electronic absorption spectroscopy was utilized to verify the compound integrity. Global fitting analysis was performed using CarpetView (Light Conversion, Vilnius, Lithuania).

**Unbiased classical molecular dynamics analysis.**
The MD system was prepared using Gaussian16 and the Antechamber program from AmberTools22.[10-12] EY geometries were first optimized at the B3LYP/6-31G* level of theory, followed by electrostatic potential calculations using the Merz-Singh-Kollman method in Gaussian16.[11] Partial charges were derived using the RESP program within AmberTools.[13,14] Starting coordinates for the MD simulations were taken from the Chai-1 predicted structures.

Each protein-ligand complex was solvated with a TIP3P water model[15] in a periodic box with a 9 Å buffer distance from the protein surface, and the system was neutralized with sodium and chloride ions. All simulations were performed in Amber18 using the ff14SB and GAFF2 force fields.[16,17] Energy minimization was performed with 1,000 steps of steepest descent followed by up to 6,000 steps of conjugate gradient. The system was heated to 310.15 K over 50 psec in the NVT ensemble using a Langevin thermostat and a 1 fsec timestep. Simulations then transitioned to the NPT ensemble, maintaining a pressure of 1 atm via a Monte Carlo barostat.[18] During the equilibration phase, heavy atoms of the protein and ligand were harmonically restrained with a force constant of 10 kcal/(mol·Å$^2$), which was gradually reduced to 0 kcal/(mol·Å$^2$) across 9 steps totaling 200 psec. Finally, an unrestrained production run was carried out under periodic boundary conditions using a 2 fsec timestep. Three independent 500 ns simulations were performed for each complex. The SHAKE algorithm[19,20] was applied to constrain hydrogen bonds. Long-range electrostatics were handled by the Particle Mesh Ewald (PME) method[21], with a 10 Å cutoff applied to short-range non-bonded electrostatic and Lennard-Jones interactions. Trajectories were analyzed using the Python packages MDAnalysis[22,23] and ProDy.[24]

**Western blot protocol.**

For Western blotting analysis, proteins were loaded on 4-12% 17-well BisTris gels (Thermo Fischer, NW04127BOX) and transferred from SDS-PAGE gels to PVDF or NC membranes (Thermo Fischer, IB24002, IB23002, IB33002, IB34002) using a dry blotting system (iBlot-2, Thermo Scientific, IB21001; iBlot-3, Thermo Scientific, IB31001). Membranes were blocked with Intercept® (TBS) blocking buffer (LI-COR, 927-60001) or 5% BSA in Tris buffered saline containing 0.1% Tween-20 (TBST, 37mM sodium chloride, 20mM Tris, 2.7mM potassium chloride, 0.05% Tween 20; pH=7.4) made in house, incubated with the primary antibodies as indicated by vendors, washed with TBST and incubated with secondary antibodies sequentially including anti-rabbit IgG Goat IR800 secondary antibody (Rockland, 926-32211), anti-rabbit IgG Goat IR680 secondary antibody (Rockland, 611-144-002), anti-rabbit IgG Goat secondary antibody peroxidase (Rockland, 611-1302), anti-mouse IgG Goat IR800 secondary antibody (Rockland, 610-145-211) and/or anti-mouse IgG Goat IR680 secondary antibody (Rockland, 610-144-002). Immunoblots images were captured by an infrared imager (LI-COR Odyssey CLx or BioRad ChemiDoc MP). Analysis was performed using ImageStudioLite.

**Mammalian cell culture and transfection.**

A549 cells and HEK293T cells were acquired from the UCSF cell culture and banking services or CellBank (National Collection of Authenticated Cell Cultures, Biological Resources Programme, CAS) and cultured in Dulbecco's Modified Eagle Medium (DMEM, Thermo Scientific, 11995073), supplemented with penicillin (50 µg/mL), streptomycin (50 µg/mL), and 10% (v/v) FBS. Transfection was performed using TransIT-PRO (Mirus Bio, fMIR 5740) or TransIT-293 (Mirus Bio, MIR 2704) according to the manufacturer's instructions.

**EYClamp labeling assay using purified proteins.**

Purified proteins with or without EY bound as indicated were incubated in a mixture with indicated photo-probe reagents (aryl-diazirine-biotin, aryl-azide-biotin or biotin-phenol) at a final concentration of 100 μM. The mixtures were then illuminated with LED for 10 min at 4 °C before 4X LDS sample buffer was added for Western Blotting analysis.

**EYClamp labeling assay in cells.**
EYClamp cellular labeling assay were performed as indicated in the workflow in **Figure 3A**. In brief, transfected A549 or HEK293T cells expressing indicated constructs were incubated at 37 °C in 5% $CO_2$ to 80% confluency and serum-starved for at least 6 hrs. They were then washed once with serum-free media and incubated with serum-free media supplemented with indicated concentration of EY for 1 h at 37 °C. Then the excessive EY were washed off with serum-free media twice and incubated with serum-free media for 15 min at 37 °C, which was subsequently repeated four times. Cells were then collected in tubes, resuspended in PBS before indicated photo-probe reagents (aryl-diazirine-biotin, aryl-azide-biotin or biotin-phenol) at a final concentration of 100 μM were added. The cells were left to incubate for 15 min at 25 °C in dark. Thoroughly mixed cells were illuminated with LED for 10 min at 4 °C and then collected for Western blotting analysis or LC-MS/MS sample preparation.

**Biotinylated protein enrichment.**
To enrich biotinylated proteins, cell pellets were resuspended in 1 mL 1X RIPA lysis buffer (Millipore Sigma, 20-188; Thermo 89900) supplemented with protease inhibitor (Protease Inhibitor Cocktail 100X, Cell Signaling Technology 5871; Roche cOmplete Protease Inhibitor Cocktail, 11697498001; Halt™ Protease Inhibitor Cocktail 100X, Thermo Scientific, 87786). After 15 min incubation on ice, cells were sonicated for 10 sec (2 sec on, 2 sec off, 20%) on a QSonica Sonicator (Q500, QSonica Sonicators, Newtown, CT) or 15 sec (2 sec on, 2 sec off, 10%) on a Scientz Sonicator (JY92-IIN, Scientz). Cell lysates were then cleared by centrifugation at 20,000 g for 10 min at 4 ˚C. Proteins were then added to 60 μL Pierce™ Streptavidin Agarose beads (Pierce, 20349) that were pre-washed with 1 mL PBS for three times and incubated for 8-16 hrs at 4 ˚C. Afterwards, supernatant was discarded and the beads were washed twice with 1 mL 1X RIPA lysis buffer, twice with 1 mL 1M NaCl in 1X PBS, and twice with 1 mL of freshly prepared 2M urea in 50 mM ammonium bicarbonate. The beads were then subjected to Western blotting analysis or LC-MS/MS analysis.

**Sample preparation for LC-MS/MS analysis.**
MS sample preparation was prepared as previously described.[2] In short, proteins on the washed beads were then digested using the Preomics iST kit in an on-bead digestion format according to the manufacturer's instructions. In brief, washed beads were suspended in 100 μL LYSE buffer provided by the Preomics iST kit and incubated at 55 ˚C for 10 min for reduction and alkylation in dark. Once the beads cooled down to room temperature, 50 μL of pre-reconstituted DIGEST were added to the beads and incubated at 37 ˚C for 3 hours with shaking. The digested peptides were then collected in the flow-through using mini Bio-Spin columns (Bio-Rad) before 100 μL of STOP solution was added and mixed using vigorous vortexing. Then the peptides were desalted using the Preomics desalting columns before they were dried under vacuum and resuspended in

15 µL solvent A (0.1% formic acid with 2% acetonitrile) for LC-MS/MS analysis. Peptide amount was monitored using a fluorometric peptide quantification kit (Thermo Scientific, 23290).

**Proteomics analysis of digested peptide samples.**

Proteomics analysis was performed as previously described.[2] In short, proteomics experiments were performed on a TimsTOF PRO (Bruker) equipped with a CaptiveSpray source and a nanoElute system. The peptides were separated on a 25 cm, ReproSil c18 1.5 µM 100 Å column (PepSep, PN. # PSC-25-150-15-UHP-nc) using a stepwise linear gradient method with water in 0.1% formic acid (solvent A) and acetonitrile with 0.1% formic acid (solvent B): 5-30% solvent B for 90 min at 0.5 µL/min, 30-35% solvent B for 10 min at 0.6 µL/min, 35-95% solvent B for 4 min at 0.5 µL/min, 95% hold for 4 min at 0.5 µL/min). Acquired data was collected in a data-dependent acquisition mode with ion mobility activated in PASEF mode. MS and MS/MS spectra were collected with m/z ranging from 100 to 1700 in positive mode.

**Analysis of proteomics dataset.**

All acquired data was searched using PEAKS online Xpro 1.6 (Bioinformatics Solutions Inc.) or FragPipe powered by MSFragger (v3.7). Spectral searches were performed using a curated FASTA-formatted dataset containing Swiss Uniprot-reviewed human proteome along with protein sequences from the designed fusion protein constructs. A precursor mass error tolerance was set to 20 ppm and a fragment mass error tolerance was set at 0.03 ppm. Peptides, ranging from 6 to 45 amino acids in length, were searched in semi-specific trypsin digest mode with a maximum of three missed cleavages. Carbamidomethylation (+57.0214 Da) on cysteines was set as a static modification while methionine oxidation (+15.9949 Da) and lysine acetylation (+42.0115 Da) were set as a variable modification. Peptides were filtered based on a false discovery rate (FDR) of 1%. Samples were normalized using total ion current (TIC) or EGFR. For p-value calculations, Welch's ANOVA was implemented in PEAKS DB software.

**Software.**

Data was analyzed and visualized using GraphPad Prism (v8.0.1) and Microsoft Excel (v16.22), in addition to software listed by each experiment. MD simulation was performed using Gaussian16 and Amber22. DNA and protein sequences were processed using SnapGene (v6.0.2). Proteomics data were analyzed by PEAKS online (Xpro 1.6) and FragPipe powered by MSFragger (v3.7). Images were made using ImageStudioLite (v5.2.5), Adobe Illustrator (v22.1) and BioRender (v2.0). Structural prediction was performed using AlphaFold3 and Chai-1 and visualized using PyMol (v3.1.6.1). Endnote (v2025) was used. Code for bioinformatics analysis performed in **Figure 4B** was made publicly available on github: https://github.com/lindseyzlin/chord-diagram/tree/main

**Statistics and Reproducibility.**

All data were derived from at least two biological replicate experiments. Statistical analyses (unpaired Student's t-tests) were performed using GraphPad Prism. Data were presented as the mean ± SD, *P≤ 0.05, **P≤ 0.01, ***P ≤ 0.001, ****P ≤ 0.0001 and n.s., not significant.
